# Genotypic and antigenic study of SARS-CoV-2 from an Indian isolate

**DOI:** 10.1101/2020.06.10.140657

**Authors:** Ruby Dhar, Akhauri Yash Sinha, Ashikh Seethy, Sri Anusha Matta, Karthikeyan Pethusamy, Trymbak Srivastava, Sunil Singh, Indrani Mukherjee, Sajib Sarkar, Rashmi Minocha, Kakali Purkayastha, Jai Bhagwan Sharma, Suman Paine, Subhradip Karmakar

## Abstract

Coronaviruses (CoVs) are one of the largest groups of positive-sense RNA virus families within the Nidovirales order, which are further classified into four genera: alpha, beta, gamma, and delta. Coronaviruses have an extensive range of natural hosts and are known to be responsible for a broad spectrum of diseases in multiple species. Severe acute respiratory syndrome coronavirus 2 (SARS-CoV-2) is the causative agent of the ongoing coronavirus disease 2019 (COVID-19) that has unleashed a global threat to public health and the economy. Coronaviruses are extensively present in birds and mammals, with horseshoe bats (*Rhinolophus affinis*), being the reservoir for the ongoing SARS-CoV-2 that seems to have resulted from a zoonotic spillover to the human host, causing respiratory infections, lung injury and Acute Respiratory Distress Syndrome(ARDS). About six coronavirus serotypes are linked with the disease in humans, namely HCoV-229E, HCoV-NL63, HCoV-OC43, HCoV-HKU1, SARS-CoV, SARS-CoV-2, and MERS-CoV. SARS-CoV-2 is the seventh CoV to infect humans. We analyzed the genome sequence of CoV-2 from isolates derived from China as well from India and encountered minute variations in their sequence. A cladogram analysis revealed the predominant strain circulating in India belongs to the A2a clad. We took one such strain (MT012098) and performed a rigorous *in-silico* genotypic and antigenic analysis to identify its relatedness to other strains. Further, we also performed a detailed prediction for B and T cell epitopes using BepiPred 2.0 server and NetCTL 1.2 server (DTU Bioinformatics), respectively. We hope this information may assist in an effective vaccine designing program against SARS-CoV-2.

## Introduction

The global outbreak of Beta (β)corona virus- is not so rare as we witnessed SARS-CoV, a member of the beta coranviradea in 2002-2003, and MERS-CoV in 2016-2018. What is, however, alarming is that the outbreak from this family of virus results in significant mortality worldwide. The striking difference between these three members, SARS-CoV-2 and SARS-CoV, MERS-CoV-2, is their contagious nature, with SAS-CoV-2 demonstrating significantly high contagiousness compared to the rest of the family members, that resulted in the global pandemic. SARS-CoV-2 and MERS-CoV-2 are both closely related to SARS-CoV-2, possibly originating from horseshoe bat reservoirs. The biological differences between these viruses are striking, with a strong genomic heterogeneity even with a high sequence homology as observed between SARS-CoV-2 and SARS-CoV-2. These sequence homogeneities are critical for their specific identification in diagnostics. The PCR-based methods play a vital role in the detection of viral infection during the early phase for clinical management by using SARS-CoV based primers. The SARS-CoV-2 virus was first sequenced by Wu et al. in Shanghai, China (Wu et al, 2020). Information derived from whole-genome sequencing as well as from partial sequences of CoV-2revealed the Genome to be29,811 nucleotides long, broken down as follows: 8,903 (29.86%) adenosines, 5,482 (18.39%) cytosines, 5,852 (19.63%) guanines, and 9,574 (32.12%) thymines that makes up the genes for the Spike (S), Nucleocapsid (N), Envelope (E) and ORF genes. The whole-genome data reveals >70% sequence homology between SARS-CoV-2 and SARS CoV and >40% sequence homology with MERS. We annotated a total of 250 genome sequences available in public databases, which includes about 90 whole-genome sequences (Hatcher et al.). This has led us to identify about 900 variant sites for the SARS-CoV-2. Genome sequencing of the SARS-CoV-2 has been done across the world. From India, two complete genome sequences deposited at NCBI with GenBank ID (MT012098, MT050493) were taken for analysis. We took MT12098 as the Genome for analyzing the antigenicity. Also, the reference genome having NCBI ID, NC_045512, was chosen as reference. The Structural proteins Spike protein(S), Nucleocapsid protein (N), Membrane Protein (M), and Envelope protein of Genome MT12098 were taken for analysis of Antigenic epitope analysis. The protein sequences between MT12098 and the Reference genome were 100% conserved. However, some variations were observed between the S protein of Reference genome and MT12098, which were otherwise 99.84% conserved. The SARS-CoV-2 Genome consists of ORF1a, ORF1b, S protein-coding region, Orf3a, E protein-coding region, M protein-coding region, ORF6, ORF7a, ORF7b, ORF8, N protein-coding region, and Orf10. The widely known structural proteins are Spike protein(S), Nucleocapsid protein (N), Membrane Protein (M), and Envelope protein (E).

### SARS CoV-2 virus phylogeneticity

Since the wide-spread effect of COVID-19 has occurred, a large number of labs have been sequencing SARS-CoV-2 genomes. Currently, more than 4000 sequenced genomes have been submitted from various parts of the world in the Global Initiative on Sharing Influenza Data (GISAID) repository (Shu et al 2017). We observed that a large number of the submitted genomes are identical, with little to no change as compared to the reference genome. We also observed some common variations in the genomes of the SARS-CoV-2. These commonly occurring variations helped scientists to classify the viral Genome. Forster et al, devised a method to classify SARS-CoV-2 based on the changes in amino acids that each of the mutations caused. The proteomes of the S clade show a Serine in the 84^th^ position of the ORF8 protein, differing from the Leucine in the reference protein. This mutation is represented as L84S, the S in the name of the clade represents the amino acid Serine. The G clade has a variation in the spike-glycoprotein, where the D at 614 of the reference protein is replaced with Glycine (G) and the mutation is named D614G. The V clade is when the Glycine of the reference protein is changed to a Valine at the 251^st^ position, ie, a G251V. This is in the NSP3 protein region (Forster et al. 2020).A large number of genomes that are submitted to GISAID are unclassified into these clades, thereby ascertaining that these clades are not extensive. According to the Nextstrain website, L84S is present in 35% genomes, the D614G mutation in 61% and G251V in 20% of genomes (Hadfield et al. 2018).

### Classification into clades based on nucleotide similarity

Another way to classify the virus is by using genomic information only(**Fig 1**). Based on this, the SARS-CoV-2 can be classified as O,A1, A1a, A2, A2a, A3, A5, B1, B2 and B4 clades. The Nextstrain website uses this classification method to draw cladograms on their website (Shu et al. 2017).The A2a type possesses a non-synonymous variant (D614G) that eases the entry of the virus into the lung cell of the host. According to the recent paper by Biswas et al, this could be the reason behind the A2a type enjoying a major advantage to infect and survive (Biswas et al. 2020).

**Figure (1A).**
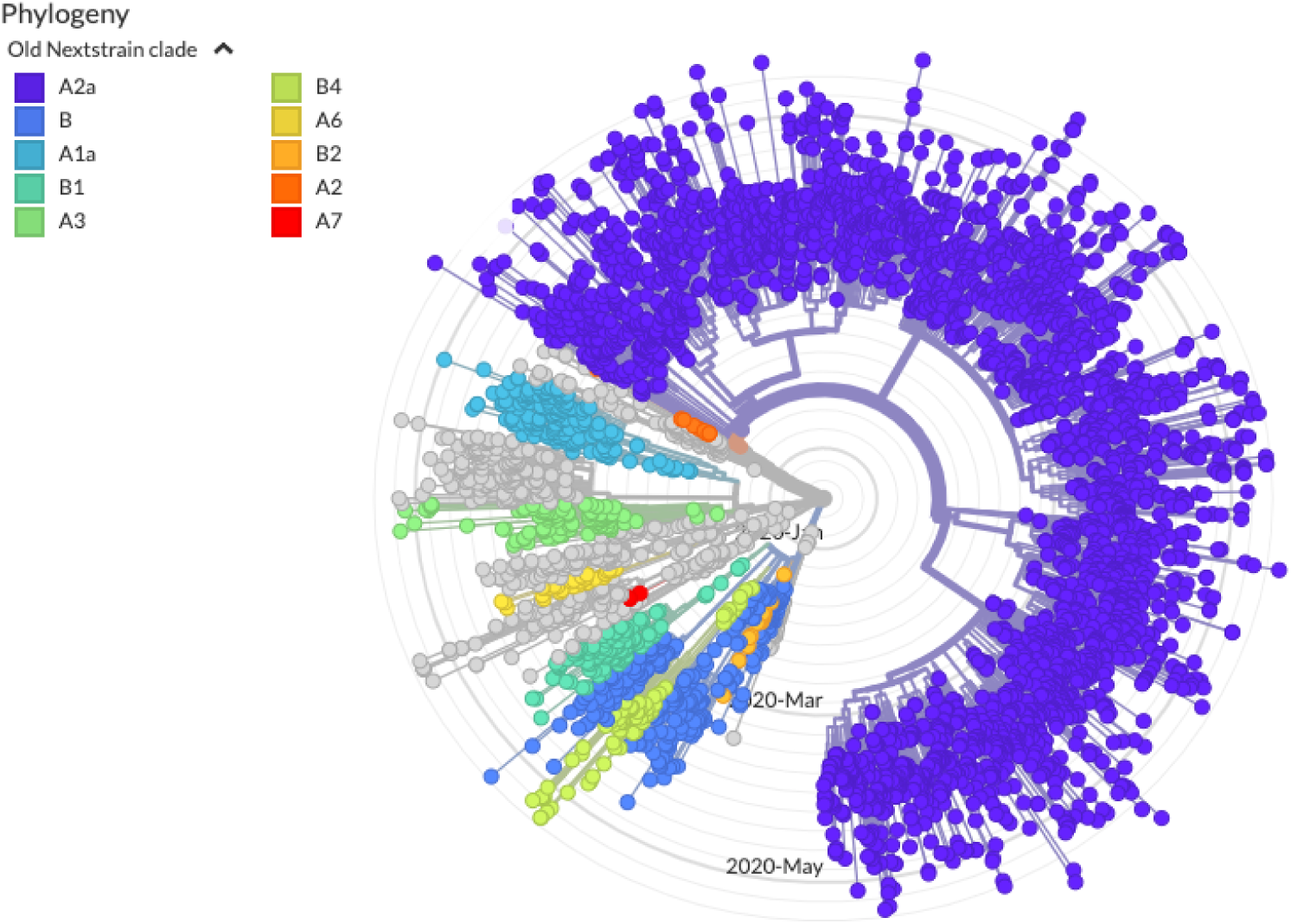
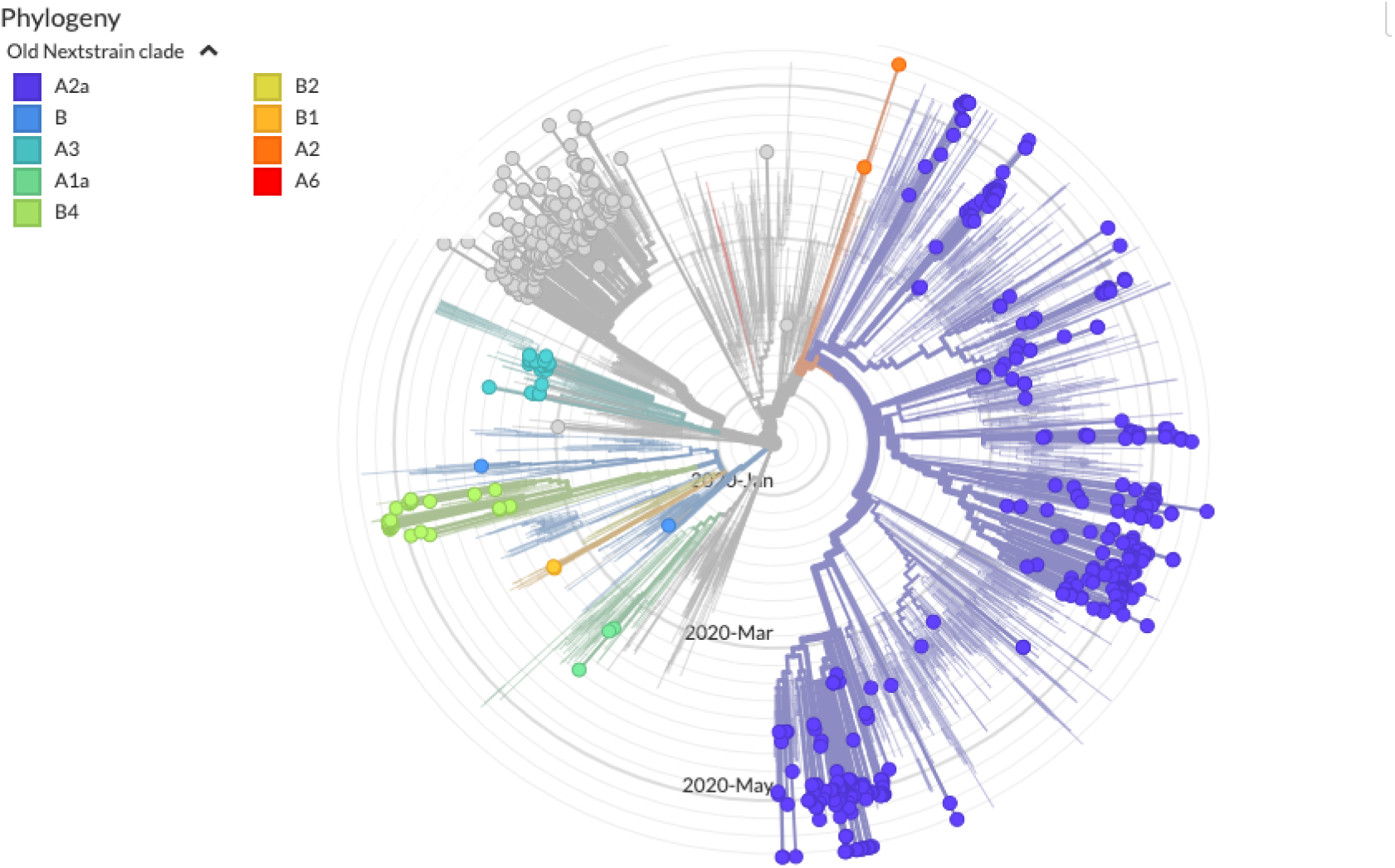
showing the clade distribution of the SARS-CoV-2 genomes sequenced globally. Most genomes lie in the A2a clade. Figure (1B) shows the clade distribution of Indian genomes.

### Indian genomes show a different perspective

So far a number of SARS-CoV-2 genomes from India have been submitted in the GISAID and NCBI websites. As seen in figure (B),the new genome strains that were published revealed that the predominant Indian strain belong to the A2a clade(60%). The remaining, close to 40% belong to the O clade (Close to the first published Genome from China)

### Phylogenetic analysis of the Indian genomes

On constructing a cladogram using NCBI’s inbuilt phylogenetic tree program, it is observed that there are wide variations in the genomes that have been submitted from India (**Fig.2**).

**Figure 2:**
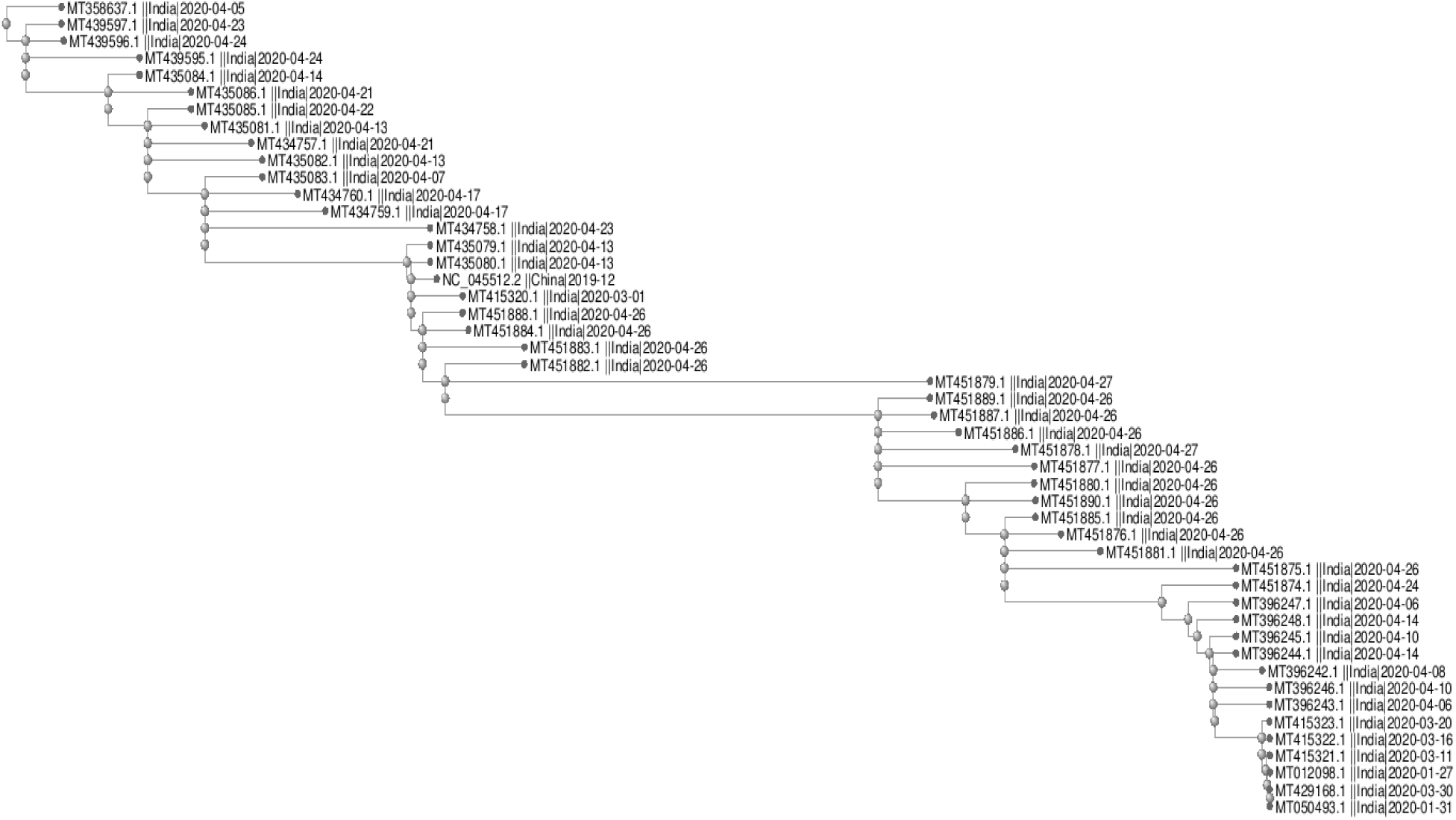
Cladogram depicting Indian genomes available on NCBI. The cladogram was constructed using distance to denote genetic mutations.

For this present study, to compare all the genomes sequenced from India, we used MT012098 as the reference genome, since this was the first whole-genome sequence that was publicly available from India. Further to note that MT012098 shows similarity to the original SARS-CoV-2 Genome published from China with a few mutations as depicted below. This Genome is categorized into the original (O) clade.

As compared to the SARS-CoV-2 from China, the Indian isolate MT012098, showed no sequence difference among the structural proteins (S,E,M and N protein). However we observe changes in the S region where there is a gap at 145 and an R407I mutation (**Fig.3**).

**Figure 3:**
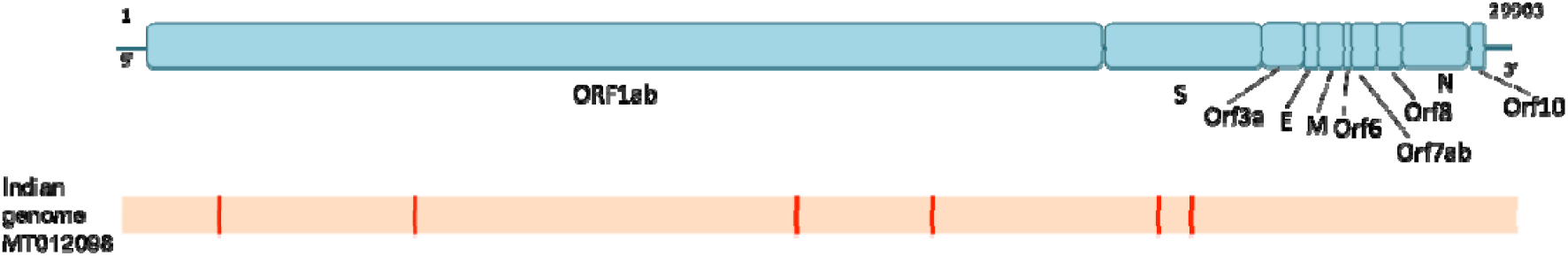
A graphical representation showing the variations of the SARS-CoV-2 Genome (in blue from an isolate in China) with MT012098 (in orange from an isolate in India) Red lines depict the variations in the Genome from the Indian isolate.

### Antigenicity of SARS-CoV-2 and possible vaccine target for SARS-CoV-2

Antigenicity can be viewed as the potential of a defined chemical structure to interact and bind to immune cell surface receptors (T cell receptors or B cell receptors (Antibodies) such as. Antigens may be, therefore be capable of eliciting an immune response due to their interaction with the immune cells. The antigens have a fundamental unit called epitopes (antigens are made of multiple epitopes) to which the Antibodies or T cell receptors can bind. Therefore, we can classify the epitopes into two broad categories, B cell epitopes and T cell epitopes. The antigenicity and immunogenicity (capacity to elicit an immune response) are therefore based on their B cell and T cell epitopes. For B cells, the epitopes could be further subdivided into (1) linear and (2) discontinuous epitopes wherein linear epitope is a continuous stretch of peptides, whereas discontinuous epitopes are distant amino acids coming near each other in space due to protein folding. Taken together, both these epitopes contribute to the overall B cell-related antigenicity/immunogenicity. The T cell epitopes are peptides or 9 to 11 or 13 to 18 amino acids, which require a special molecule called Major histocompatibility complex to bind to them and be presented on the cell surface through a classical antigen receptor cells (APC). The T-cell receptor then recognizes both MHC and the peptide presented for binding and eliciting the immune response. The peptide is presented by broadly two kinds of MHC molecules. MHC class I and MHC class II. Wherein, peptide presented with MHC class I (8 to 10 amino acids) elicit CD8+, Cytotoxic T cell lymphocyte (CTL) response. and CD4+ response is elicited by peptide presented by MHC class II.

The cytotoxic T lymphocytes are responsible for the execution of apoptotic cell death of the infected cell through the release of cytotoxic molecules like cytokines and interferons On the other hand; the CD4+ T cell is majorly responsible for assisting the humoral immunity (Inducing the B cell response).

## Material and Methods

Complete Genome of SARS CoV-2 strains Reference genome NCBI ID, NC_045512, and Indian genome GenBank ID, MT012098, were taken from NCBI website.Blast: **Blast** P of NCBI was used for Structural protein comparisons between reference strain NC_045512 and Indian Strain MT012098

**Figure 4:**
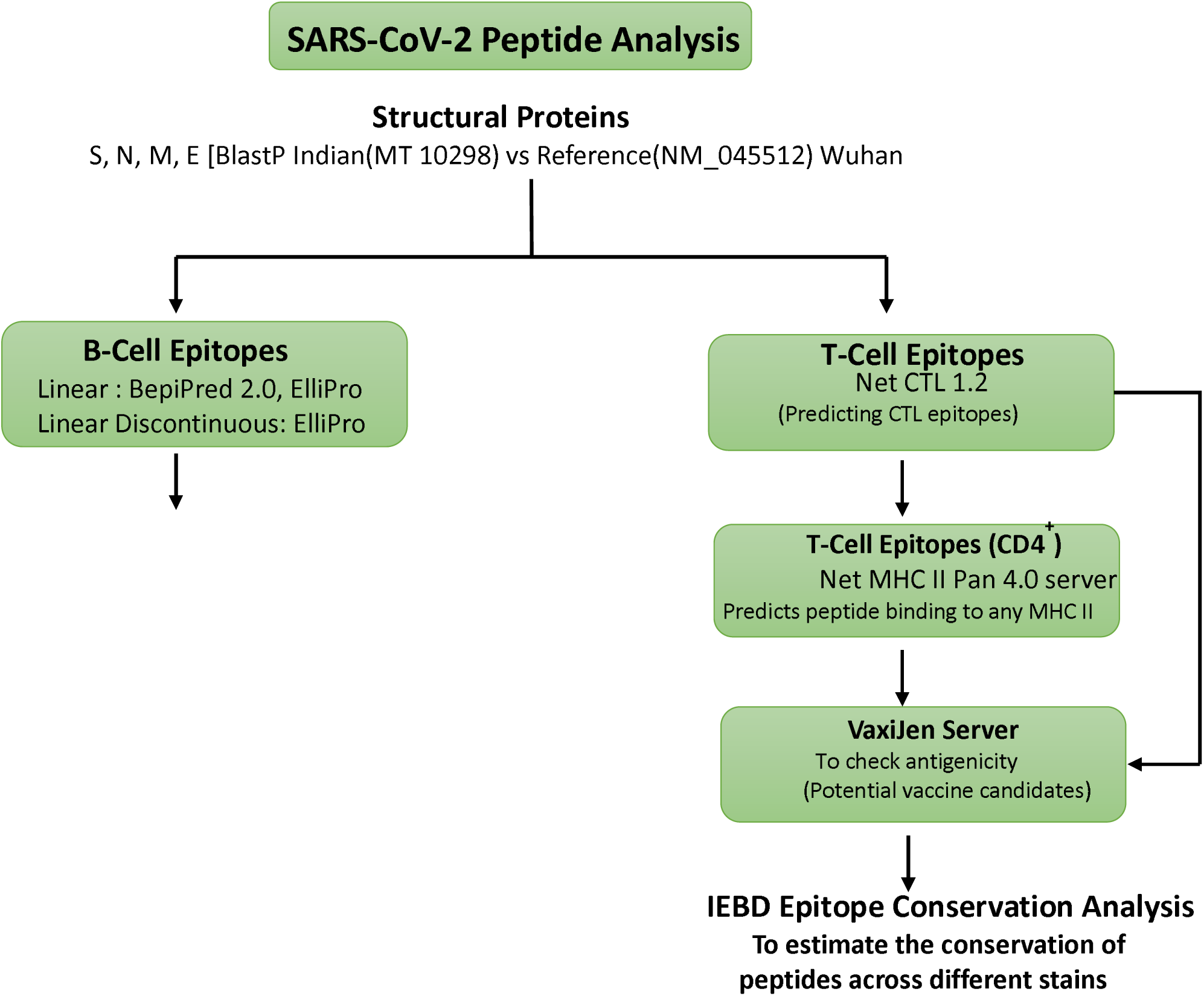
Summary of the workflow for antigenic peptide prediction.

### B cell Epitope prediction

For B cell linear epitopes Bepipred version 2 algorithm was used. The software utilizes the random forest algorithm, which is trained on epitopes annotated from the antibody-antigen protein structure (Jespersen et al., 2017). The Bepipred version 2 is available in Immune Epitope Data Base Analysis resources (http://tools.iedb.org/bcell/). Further, ElliPro (http://tools.iedb.org/ellipro/) was used for the analysis of both linear and discontinuous analysis(Ponomarenko et al., 2008). ElliPro (Ellipsoid and Protrusion), is a web-tool that implements a modified version of Thornton’s method(Thorton et al. 1986).

### T cell Epitope Prediction

#### CTL epitopes (MHC Class I associated peptides)

For T cell epitopes MHC class I, analysis NetCTL1.2(http://www.cbs.dtu.dk/services/NetCTL/) was used, and eight supertypes were analyzed. These were HLA A1, A2, A3, A24, B7, B27,B58 and B62. NetCTL1.2 utilizes the processes required for antigen presentation for its algorithm and calculation of score (Larsen et al., 2007). Briefly, Scores are based on

(1) MHC binding affinity, (2) Proteasomal cleavage, (3) Tap association (4) Combine score

The default cut off for the Combine score is 0.75, which was considered.

#### CD4^+^ epitopes (MHC class II-associated peptides)

For CD4^+^ epitopes, NetMHCIIpan 4.0 (http://www.cbs.dtu.dk/services/NetMHCIIpan/(Reynisson et al., 2020) was used for HLADP, HLA DQ and HLA DR alleles. Peptides with a rank percentage of less than 2%, which is the default threshold of strong binders, only were considered.

#### Antigenicity analysis

VaxiJen Version 2.0 (http://www.ddg-pharmfac.net/vaxijen/VaxiJen/VaxiJen.html) was used for measuring antigenicity and for alignment independent prediction of protective antigens. It uses alignment-free prediction based on the physicochemical properties. It utilizes autocross covariance(ACC) for its algorithm. The algorithm is trained on 100 antigens and 100 non-antigens of bacterial viral and tumor proteins(Doytchinova and Flower, 2007).

#### Conservancy analysis

Conservancy analysis for the peptide was performed using IEDB epitope conservancy analysis tool http://tools.iedb.org/conservancy/(Bui et al., 2007).

## Results

### Blast

Blast P for S,N M, and E protein were performed between the reference Genome and NC_045512 (Severe acute respiratory syndrome coronavirus 2 isolate Wuhan-Hu-1, complete Genome)and the Indian isolate MT012098. For N, M, and E protein, 100% identity was observed while for S protein, 99.84% identity was with the variations above mentioned. Due to these variations, S protein of the reference strain was separately used for analyzing the antigenic peptides for B cell and T cell epitopes. Similarly, S, M, N, and E protein of the Indian strain were also utilized for B and T cell peptide analysis.

### Prediction of B Cell Epitopes

For analyzing the linear epitopes, IEDB BepiPred linear epitope prediction tool was used (http://tools.iedb.org/bcell/)(Jespersen et al., 2017). The threshold cut off was taken as 0.5, which is also the default cut-off. A total of 33 B cell epitope peptides were given as output for reference MT012098 S protein, while11,6, and 2 for N, M, and E protein were observed, respectively (Table 1a, 1b, 1c, 1d). The variation in B cell linear epitope antigenic peptides of S protein reference strain compared to the Indian strain is shown (**Fig 5A-5C**).

**Fig 5A:**
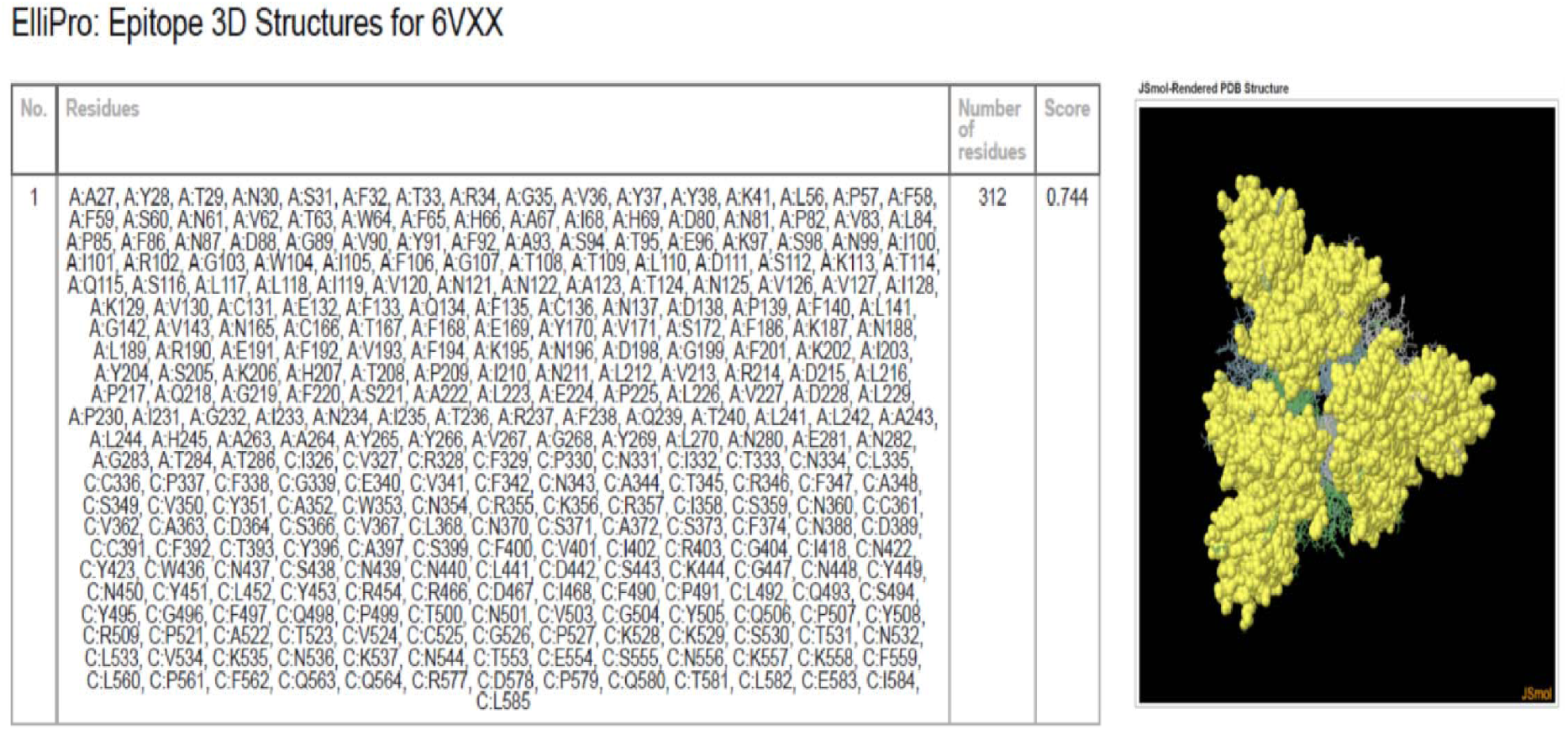
Describes the Discontinuous epitopes of S protein. A and C represents Chain A and C for S protein. The yellow region represents the antigenic area.

**Fig.5B:**
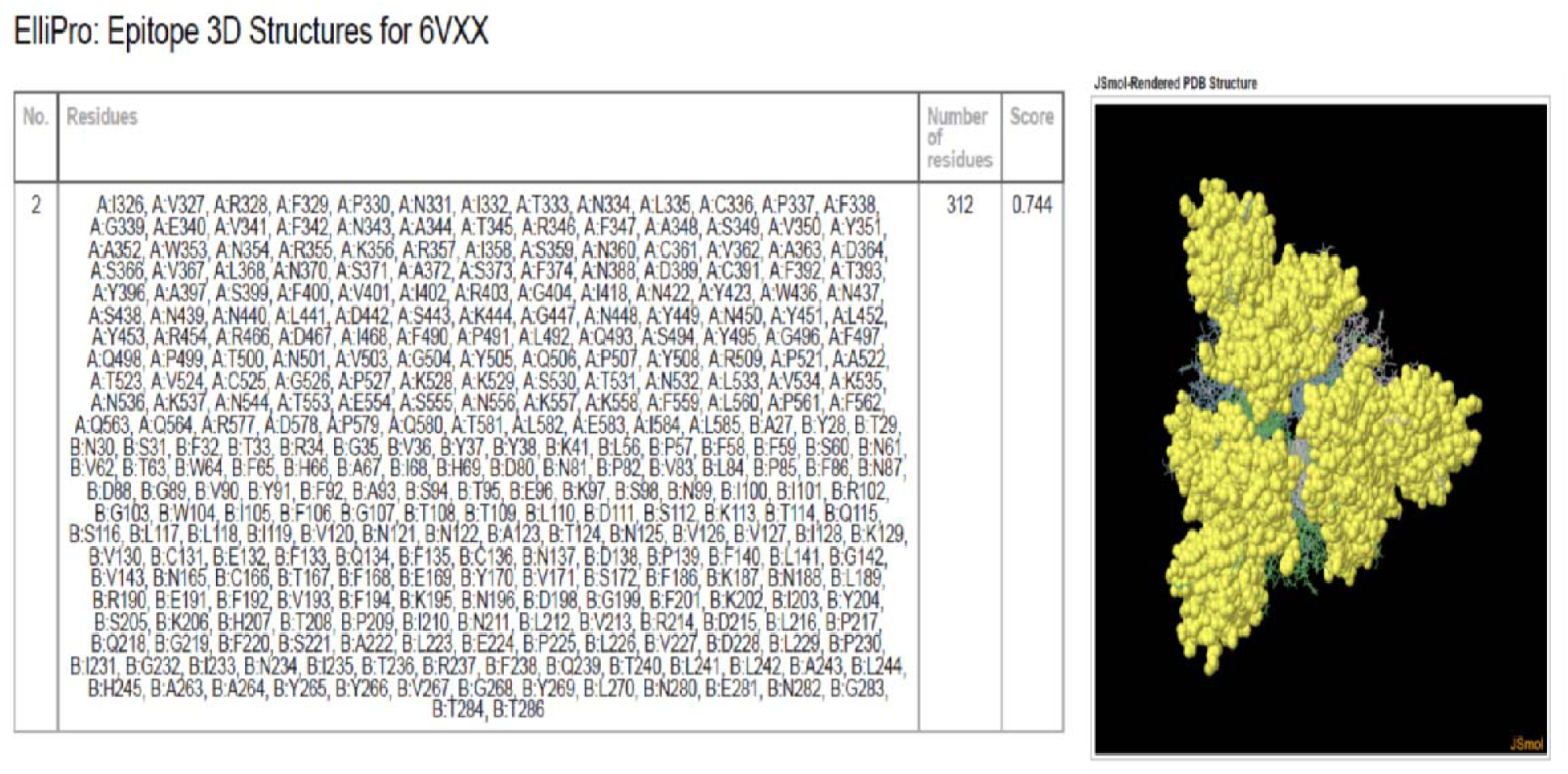
Describes the Discontinuous epitopes of S protein. A and B represents Chain A and C for S protein. The Yellow region represents the antigenic area.

**Fig.5C :**
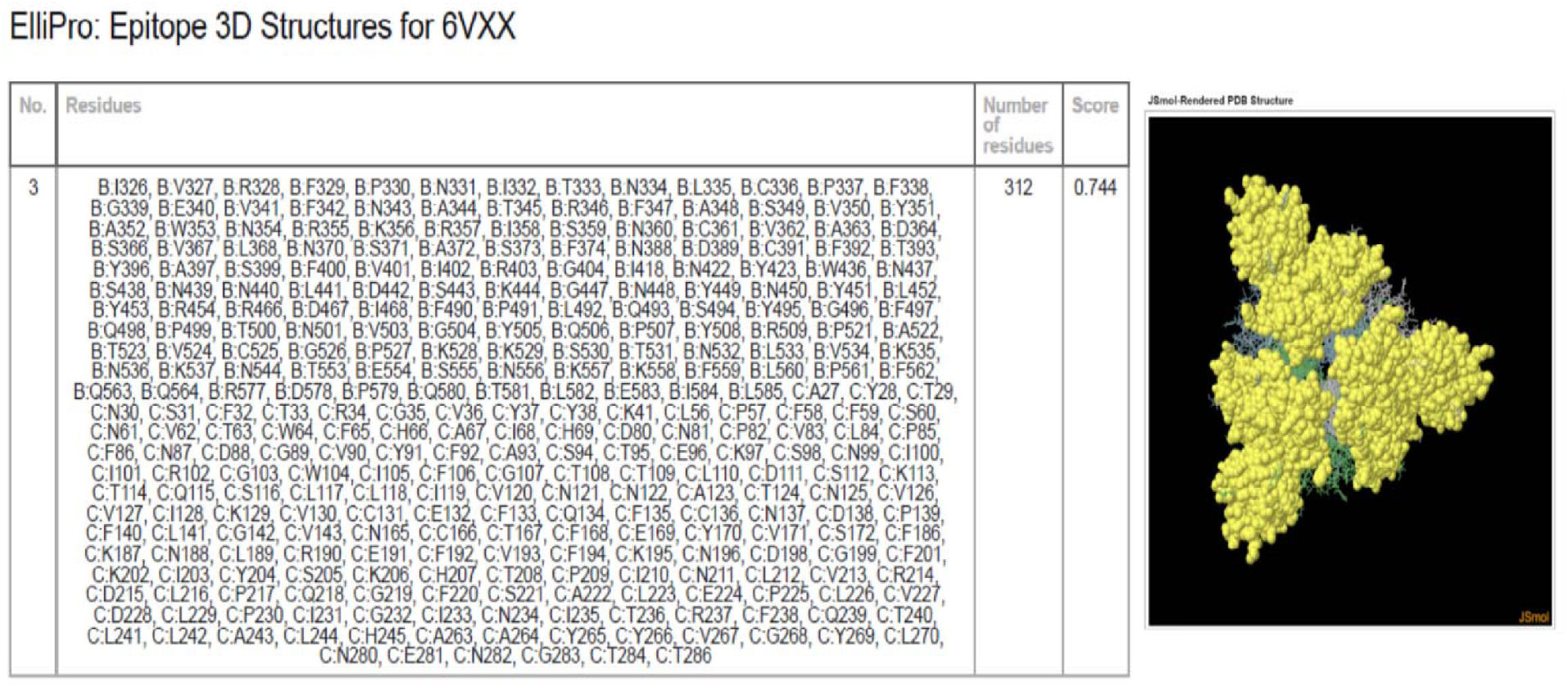
Describes the Discontinuous epitopes of S protein. B and C represents Chain B and C chains for S protein. The Yellow region represents the antigenic area.

**Fig. 5D:**
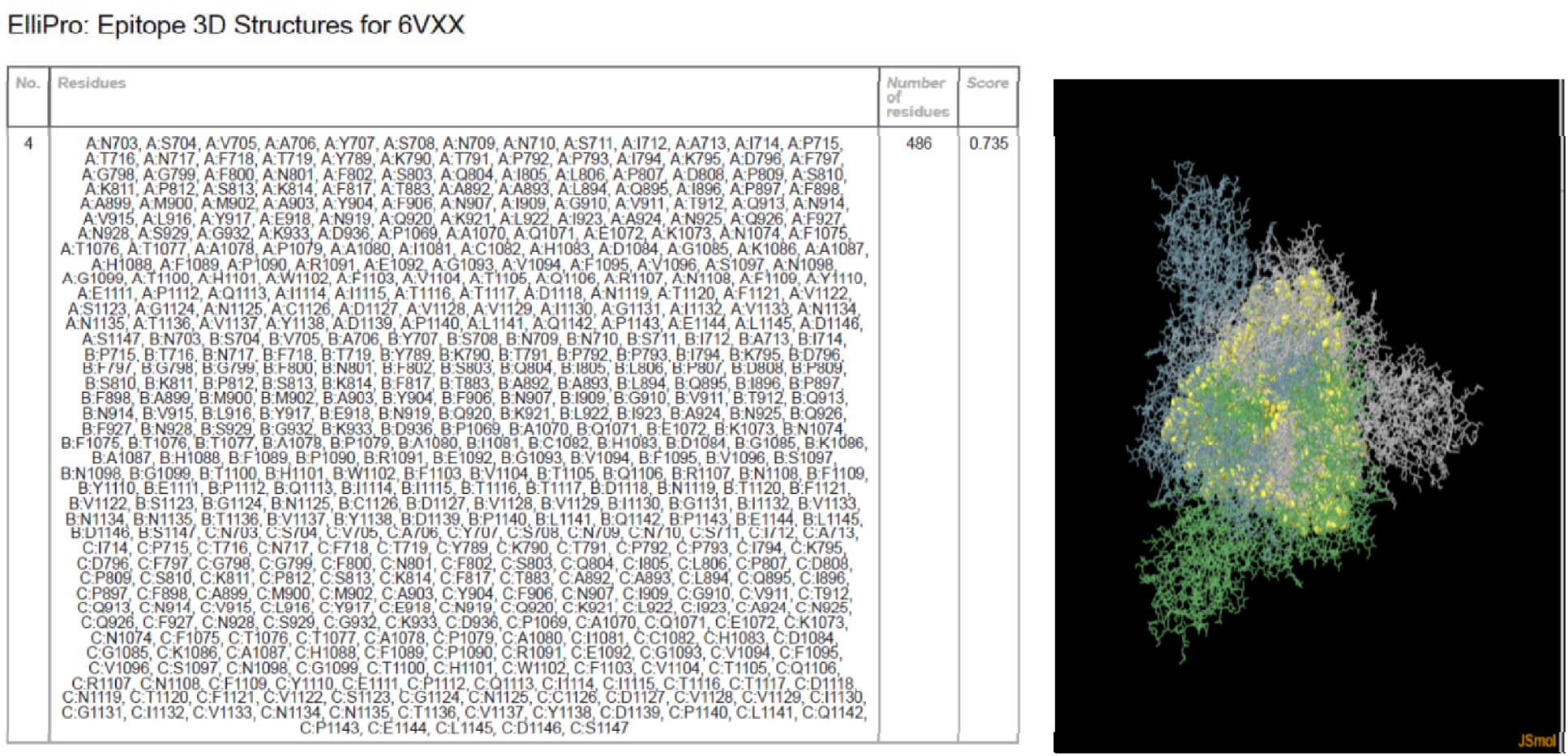
Describes the Discontinuous epitopes of S protein. A, B and C represents Chain A,B and C chains for S protein. The Yellow region represents the antigenic area.

For discontinuous epitopes, the IEDB Ellipro algorithm was used with the threshold cut-off based on the protrusion index (P.I.) score is 0.5 (Ponomarenko et al., 2008). However, we had taken the cut-off of 0.7 as a greater score means greater solvent accessibility(Adhikari et al., 2018; Ponomarenko et al., 2008). The maximum distance between the amino acids in space was taken as 6 angstrom, which is the default cut-off. The S protein and N protein alpha chain of only the reference genome was analyzed as the Protein Data Base (PDB) I.D., and the structure of the protein in PDB is required for the algorithms analyzing discontinuous B cell epitopes. Minimal variation was observed for the P.I. score above 0.7. The maximum was 0.744 and minimum of 0.735 (**Fig 5 A-D**). Four different epitopes with P.I. score of more than 0.7 was observed. For the N protein alpha chain, only one epitope was observed, having a score greater than 0.7.

### T cell epitope prediction

For analyzing MHC, class I represented T cell epitopes, NetCTL version 1.2 was used. The advantage of this algorithm, in addition to utilizing the process required for MHC class I representation of T cell peptides, is that it has a pool of MHC haplotypes grouped into MHC supertypes(Dos Santos Francisco et al., 2015; Sidney et al., 2008; Zhao et al., 2017). NetCTL 1.2 has haplotype grouped into12 MHC supertypes, which are HLA A1, A2, A3, A24, A26, B7, B8, B27, B39, B44, B58, and B62. Among these, the more common HLA A1, A2, A3, A24, B27, B44, and B62 were chosen. The default cut-off of 0.75 for the combined score was determined. The higher score has improved specificity but reduced sensitivity while the lower score has more sensitivity but with reduced specificity. For MT012098, S protein, a total of 314 potential MHC Class I associated T-cell peptides were obtained for all the 8 MHC supertypes(data not shown). Similarly, 75,184 and 34 peptides were obtained for N, M, and E proteins of MT012098, respectively. Each of these peptides was further subjected to antigenicity analysis using the VaxiJen server 2.0. The default cut-off score for potential antigens is 0.4; however, due to a large number of hits, we increased the threshold to 0.7. A super peptide, which was having a score greater than 0.7 were only considered. These peptides had a high probability of associating with the surface MHC class I molecules and have high probable antigenicity and therefore has a strong vaccine potential. The peptides for S protein having a cut-off score over 0.7 have been depicted in Figures 5A and 5B. A total of 98 potential super peptides was obtained with 44 belonging to HLA A(4 supertypes) haplotype while 54 for HLA B (4 supertypes) haplotype for MT012098. Notably, for N,M and E 26, 40, and 14 supertypes for all the above-aforementioned supertypes were obtained for MT012098 (**Table 2C,2D, 2E**).

Only three potential T cell peptides derived from S protein of the reference genome were different form the S protein of MT012098. These peptides of the S protein of the reference genome were ^142^’GVYYHKNNK’^150, 144^’YYHKNNKSW’^152^and ^408^’RQIAPGQTG’^416^. Among the exclusive peptides, ^142^’GVYYHKNNK’^150^, and ^408^ ‘RQIAPGQTG’ ^416^ were considered as super peptides as they had a score of more than 0.7 for antigenicity in VaxiJen Server 2.0. The peptide ^144^’YYHKNNKSW’^152^, although had a higher score than the default cut-off score of 0.4, was not considered as Super peptide because its antigenicity score was less than 0.7

### T cell epitope CTL (MHCI epitope prediction)

For analyzingMHC, class, I represented T cell epitopes, NetCTL version 1.2 was used. The advantage of this algorithm, in addition to utilizing the process required for MHC class I representation of T cell peptides, is that it has a pool of MHC haplotypes grouped into MHC supertypes(Dos Santos Francisco et al., 2015; Sidney et al., 2008; Zhao et al., 2017). NetCTL 1.2 has haplotype grouped into12 MHC supertypes, which are HLA A1, A2, A3, A24, A26, B7, B8, B27, B39, B44, B58, and B62. Among these, the more common HLA A1, A2, A3, A24, B27, B44, and B62 were chosen. The default cut-off of 0.75 for the combined score was determined. The higher score has improved specificity but reduced sensitivity while the lower score has more sensitivity but with reduced specificity. For MT012098, S protein, a total of 314 potential MHC Class I associated T-cell peptides were obtained for all the 8 MHC supertypes(data not shown). Similarly, 75,184 and 34 peptides were obtained for N, M, and E proteins of MT012098, respectively. Each of these peptides was further subjected to antigenicity analysis using the VaxiJen server 2.0. The default cut-off score for potential antigens is 0.4; however, due to a large number of hits, we increased the threshold to 0.7. A super peptide, which was having a score greater than 0.7, was only considered. These peptides had a high probability of associating with the surface MHC class I molecules and have high probable antigenicity and therefore has a strong vaccine potential. The peptides for S protein having a cut-off score over 0.7 have been depicted in **Fig 5A and 5B**. A total of 98 potential super peptides was obtained with 44 belonging to HLA A(4 supertypes) haplotype while 54 for HLA B (4 supertypes) haplotype for MT012098. Notably, for N, M and E 26, 40, and 14 supertypes for all the above-aforementioned supertypes were obtained for MT012098 (**Table 2C,2D, 2E**).

We identified three potential T cell peptides for S protein for Indian strain MT012098 from the reference genome. The exclusive peptides of S protein of the reference genome were ^142^’GVYYHKNNK’^150, 144^’YYHKNNKSW’^152,^ and 408’RQIAPGQTG’^416^. Among the exclusive peptides, ^142^’GVYYHKNNK’^150^, and 408’RQIAPGQTG’416 were considered as super peptides as they had a score of more than 0.7 for antigenicity in VaxiJen Server 2.0. The peptide ^408^’RQIAPGQTG’^416^, although had a higher score than the default cut-off score of 0.4, was not considered as Super peptide because its antigenicity score was less than 0.7.

### T cell epitope CD4^+^ (MHC II epitope prediction)

Similar to T cells, we performed CTL MHC class I and T cell CD4^+^ MHC class II analysis. NetMHCIIpan 4.0 was used to predict the potential MHC II-associated peptides for the CD4^+^ T cell response. As the human MHC types comprise of HLA DP, HLA DQ, and HLA DR types. We had chosen the twenty most common HLA DP (n=5), HLA DQ (n=5), and HLA DR (n=10) haplotypes present in the world or India for the analysis(Shankarkumar, 2010; Shen et al., 2018). HLA DR is more abundant in the human population; therefore, more numbers of HLA DR were considered. The HLA DR MHC type includedDRB10101, DRB1_0301, DRB1_401, DRB1_404, DRB1_0701, DRB1_0802, DRB1_0901, DRB1_1001, DRB1_1501, DRB1_1508. The other major MHC types HLA DP and HLA DQ included DPA10103-DPB10201, DPA10103-DPB10301, DPA10103-DPB10401, DPA10103-DPB10501, DPA10201-DPB10101 and HLA-DQA10101-DQB10501, HLA-DQA10102-DQB10502, HLA-DQA10102-DQB10602, HLA-DQA10103-DQB10603, HLA-DQA10104-DQB10503 respectively. The predicted MHC II-associated peptide was obtained from Net MHCII pan 4.0 under stringent conditions.NetMHC 11 Pan-4.0 Server predicts peptide binding to any MHC II molecule of known sequence using Artificial Neural Network (ANN). Only those who had rank percentage less than 2%, which is the default threshold of the strong binder, was considered. The weak binders were eliminated. A total of 124 potential peptides was obtained for the different sub haplotypes of HLA DR and 73 and 56for different sub haplotypes of HLA DP and HLA DR, respectively. These peptides were further subjected to antigenicity analysis similar to MHC class I CTL associated peptides with the threshold of 0.7 to obtain super peptides. A total of 23 super peptides associated with different HLA DR haplotypes were obtained (**Table 3A**), while six peptides and eight peptides were obtained for HLA DP and HLA DQ, respectively (**Table 3B and Table 3C**).

Comparison between the antigenic peptides from S protein of reference genome NC_045512 and Indian genome MT012098 showed that ^404^DEVIQIAPGQTGKIA^41^ and ^398^SFVIRGDEVIQIAPG^412^ were unique to Indian S protein (MT012098 genome) while ^137^NDPFLGVYYHKNNKS^151^ and ^404^GDEVRQIAPGQTGKI^418^ were obtained from S protein of Reference genome NC_045512 and not from the Indian S protein MT012098 genome.

In addition to the S protein, the antigenic potential of N protein of MT012098 genome with respect to the MHC class II association was also analyzed. As previously shown in the paper, there is a100% match for the blast result for N protein of reference (NC_045512) with the N protein derived from the Indian (MT012098) isolate genome. No variation in antigenicity between them would, therefore, be observed. The N protein analysis included all the MHC II types, which were also used for the S protein analysis. A total of 5 peptides were observed to be super peptides across the MHC types. The peptide ^328^GTWLTYTGAIKLDDK^342^ was found to be the most dominant N protein antigenic super peptide, which displayed polymorphism for various MHC types; these included DRB1_0101, DRB1_0701, DRB1_0901, DRB1_1508, and DPA10201-DPB10101 (**Table 4A**).

### Conservation analysis

The B and T cell epitopes were analyzed for their conservation across different genomes of SARS CoV-2. Fourteen different genomes across the globe were compared with each antigenic peptide these include Reference Genome (NC_045512), Genomes sequenced for India (MT012098, MT050493). Genome sequenced for USA (MN985325, MT300186), Genome sequence from Italy (MT066156, MT077125), Genome sequenced from Greece (MT328032), Spain (MT292570), Australia (MT007544), Taiwan (MT066175), Japan (LC534419), Brazil(MT126808), and Iran (MT320891). For B cell epitopes for MT012098, S protein, 34 out of 36 peptides were 100% conserved while peptides ^94^’STEKSN’99 and^368^’ YNSASFSTFKCYGVSPTKLNDLCFT’ ^392^ were only 7.2% conserved. For N protein also, 9 out of 11 peptides were 100% conserved while ^165^’TTLPKGFYAEGSRGGSQASSRSSSRSRNSSRNSTPGSSRGTSPARMAGNGGD’^216^ and 276’RRGPEQTQGNFGDQELIRQGTDYK’^299^ were 93.86% conserved (**Table 1A** and **Table 1B**). All the potential B cell peptides for M and E proteins were 100% conserved (**Table 1C** and **Table 1D**). For the B cell epitopes, peptides that were exclusive to the reference genome, 6 out of 8 were 100% conserved. This suggests an interesting scenario that this sequence, though was present in all the different genomes, possibly didn’t emerge as potential antigenic peptides, probably due to variations in the neighboring amino acids. This suggests that neighboring amino acids play an important role in determining antigenicity. The peptides which were not 100% conserved were ^138^ ‘DPFLGVYYHKNNKSWME’ ^154^and ^404^ ‘GDEVRQIAPGQTGKIADYNYKLP’ ^426^(**Table 1E**).

The T cell CTL Super peptides besides being highly antigenic were also highly conserved. For MT012098, S protein 97 out of 98 potential super peptides were 100% conserved while ^239^ ‘TLLALHRS Y’ ^247^is 92.23% conserved (**Table 2A**). The Super peptides M, N, and E protein of MT012098 showed 100% conservancy.

The two exclusive super peptides of reference genome ^142^’GVYYHKNNK’^150, 408^’RQIAPGQTG’^416^ were 92.86% conserved. For the T cell CD4^+^ MHC class II peptides, 21 of the 23 super peptides for HLA-DR MHC types were 100% conserved while the remaining two^404^DEVIQIAPGQTGKIA^418^,^398^SFVIRGDEVIQIAPG^412^, were unique peptides with 7.69% conservation. As aforementioned, these two peptides were also not observed in the S protein of reference genomeNC_045512. However, these two peptides are highly antigenic. For HLA-DP 5 out of 6 peptides were 100% conserved while one peptide ^233^NITRFQTLLALHRSY^257^ showed 92.86% conservation matching with 13 out of 14 genomes. For HLA-DQ, however, all the eight peptides were 100% conserved (**Table3 A, B, C**). Further, ^137^NDPFLGVYYHKNNKS^151^ and ^404^GDEVRQIAPGQTGKI^418^, which were obtained from S protein of Reference genome NC_045512 and not present in Indian genome MT012098, showed 92.86% conservation, wherein it was conserved with the S protein from 13 genomes mentioned in the paper exception the Indian genome MT012098.Besides the analysis S protein associated with MHC class II, N protein analysis of the Indian Genome MT012098revealed that all the five super peptides were 100% conserved (**Table 4A**).

## Conclusion and Future Directions

SARS-CoV-2 is a contagious and highly pathogenic virus associated with adverse respiratory pathology. It is a positive single-stranded RNA virus. COVID-19 caused due to this virus is transmitted to and among humans by direct contact, droplet, and airborne routes. It is believed that this virus came originated from a zoonotic reservoir by cross-species spillover. This virus was named due to its high homology (∼80-85%) to SARS-CoV, which caused acute respiratory distress syndrome (ARDS). The mean incubation period is approximately five 1days. The confirmed case fatality ratio, or CFR (the total number of deaths divided by the total number of confirmed cases at one point in time), is approx 3-3.5%. Within China, the confirmed CFR, as reported by the Chinese Center for Disease Control and Prevention, is 2.3%, though it could be higher in different populations. The primary problem of this disease results in dysregulation of cytokines/chemokines, recruitment of immune cells with direct virus-mediated cytopathic effects. Patients with severe diseases were reported to have increased plasma concentrations of proinflammatory cytokines, including interleukin (IL)-6, IL-10, granulocyte-colony stimulating factor (G-CSF), monocyte chemoattractant protein 1 (MCP1), macrophage inflammatory protein (MIP)1α, tumor necrosis factor (TNF)-α along with an associated lymphopenia. Using comprehensive genomic and proteomic tools, we identified the different variants of SARS-CoV-2 in circulation within the world community. We focused our analysis on the Indian stain MT012098 and identified unique T and B cell epitopes that have high antigenicity and could serve as a potential vaccine candidate.

### Complex Immunobiology and Interaction

Histopathological evidence reveals that upon SARS-CoV-2, infection in humans exhibits nonspecific inflammatory responses, including edema and inflammatory cell infiltration along with severe exfoliation of alveolar epithelial cells in an organized (Fu et al. 2020). In addition, SARS-CoV-2 infection was found to cause pyroptosis in macrophages and lymphocytes (Yang et al. <u>2020</u>) Derangement of pulmonary arteriolar walls may be an indicator of the crucial inflammatory response that contributes significantly to disease progression or severity of Beta corona family virus, including SARS, MERS, and SARS-CoV-2. At autopsy of SARS-CoV-2 cases reveals the lymphoid depletion in lymph nodes(Jiang Gu. et al. 2005). Clinical features like lymphopenia and increasing viral load in the first few days of the disease strongly propose the evasion of the immune system by SARS-CoV-2. Though the data is limited, the host immune response to Beta coronavirus family, including SARS-CoV-2, is a complex phenomenon with a multigenic network that yet to explore. Initially, the involvement of ACE on SARS-CoV-2 infection and its facilitation into the host cell, the complex immune phenomenon upon antigenic presentation through Pathogen associated molecular pattern (PAMP) with the aid of TLRs family and its cytokines and chemokines secretion may play a potential role for a varied range of disease prognosis(Frieman M et al. 2008, Tay MZ et al. 2020)

### TLRs against β-coronavirus

It has been well established that exposure to lipids, peptides, and nucleic acids derived from pathogenic agents (bacterial, viral, parasite, and fungal) are recognized by host Toll-like receptors (TLRs) leading to their activation(Van Meer et al. 2008; Gay N et al. 2014). The binding of antiviral components to TLRs activate downstream signaling pathways to release cytokines as a primary inducer for inflammation and promotes secondary anti-inflammatory mechanisms through PAMPs(Pathogen activated molecular patterns). Recognition of viral glycoproteins by TLR2 and TLR4 are well established, and studies demonstrated that TLR2/6 heterodimers activate the innate immune response(Geng Li et al. 2020). It has been observed that although the TLR1/2 heterodimers recognize viral glycoproteins, the potential involvement of respiratory virus infection is unclear. Usually, TLRs recognize viral PAMPs through common adaptor molecules, including MyD88, MAL, TRAM, and TRIF(Kawai et al. 2010). The TLR adaptor molecules signal through the IKK□/TBK1 complex and the IKKα/IKKβ/IKKγ complex similarly may also involve IRAK-1/IRAK4/TRAF6 complex to activate the transcription factors including IRF3, IRF7, and NF-κBthat ultimately activate IFNs and produces proinflammatory cytokines(Pobezinskaya et al. 2008, Ermolaeva MA et al. 2008). Previous in vivo study postulates that TLRs signaling programmed by MyD88 adaptor protein and in mice model study reveals the potential involvement of TLR3/TLR4 on SARS-CoV infection(Zheng et al. 2020, Perales et al 2013). The study documented that transgenic mice deficient in the TLR3/TLR4 adaptor TRIF are highly susceptible to SARS-CoV infection, showing increased weight loss, mortality, reduced lung function, increased lung pathology, and higher viral titers. TLRs potentially involve differentiating between “self” and “nonself” to the host immune system. TLRs, particularly TLR3/TLR4, seem to act as vaccine adjuvants or antiviral molecules(Akira et al. 2006). The emergence of severe contagious β-coronaviruses, including SARS-CoV-2, SARS-CoV, and MERS-CoV, accompanied a significant mortality ratio in human populations across the globe(Astuti and Ysrafil 2020). In spite of the pandemic nature of the β-coronavirus family, the therapeutics or proper intervention strategy is yet to formulate. Previous in-vivo and animal model data postulate that TLR signaling through the mediators, including TRIF and MyD88-driven adaptor protein(Kawasaki et al. 2014), may able to maintain the homeostasis of pro and anti-inflammatory to perform effective antiviral defense mechanism against β-coronaviruses family viruses including SARS-CoV-2 and SARS-CoV. Previous data insists us to postulate that TLR antagonists may be crucial in coronavirus-specific vaccine and antiviral strategies(Nilsen et al. 2015).

### Chemokines and dendritic cells

Studies demonstrated the potential role of dendritic cells (D.C.s) as an antigen-presenting cell for several RNA viruses. Genomic characterization of the SARS-CoV-2 potential N-linked glycosylation and surface spike (S) protein is of high-mannose structure that may be crucial for binding of SARS-CoV-2 and entry into the D.C.s. The mechanism may be mediated CD206, CD209, CD207, and CD205. It has been also postulated the unique adaptive mechanism for β-coronaviruses family viruses, including SARS-CoV-2, MERS, and SARS-CoV-2, to escape host immune mechanisms. However, in-vitro studies demonstrated the incomplete replication of SARS-CoV-2, where data for SARS-CoV-2 is essential to understand the phenomenon. Altered homeostasis of antiviral pro and anti-inflammatory has been postulated for SARS-CoV-2 and SARS-CoV. D.C.s signal to ‘nonself’ for the adaptive immune response to polarize T-helper (Th) cells toward the Th1/Th2 subsets. In-vitro studies also reveal that cytokines include IFN-α, IFN-β, IFN-γ, and IL-12p40 reduce 3d expression on D.C.s and upregulation of proinflammatory cytokines TNF-α and IL-6. The chemotactic messengers, including chemokines, are potential candidates for leukocyte recruitment upon single-stranded RNA virus infection. The up-regulation of chemokines has been associated with viral diseases(Glass et al. 2003). In concordance with the detection of high-plasma concentration of chemokines in patients with β-coronavirus postulated at least for SARS-CoV-2 but need to explore for SARS-CoV-2. Mice model study also reveals the surge of chemokines in the lungs on SARS-CoV–2 infection(Shi-Hui-Sun et al. 2020; Jason et al. 2008). Elevation level was highest for inflammatory chemokines (macrophage inflammatory protein (MIP)-1α/CCL3, secreted (RANTES)/CCL-5, interferon-inducible protein (IP-10)/CXCL10 and monocyte chemotactic protein (MCP)-1/CCL2 upon SARS-Co-V infection and it may be true for other β-coronaviruses family viruses including SARS-CoV-2(Coperchini F, 2020; Zang et al. 2020). We postulated that this lack of antiviral cytokine response against a background of intense chemokine upregulation could represent a mechanism of immune evasion by SARS-CoV-2; however, the genomic heterogeneity of S and N protein between SARS-CoV-2 and CARS-CoV-2 is essentially required to understand the immune escape mechanism.

Angiotensin-converting enzyme 2 (ACE2) acts as a cell receptor that facilitates viral entry to the host cell(Bourgonje et al. 2020). The recent tissue-specific expressional analysis revealed that *ACE2* expression levels were the highest in the intestine, kidneys, heart, and adipose tissue, and moderate for lungs, liver, bladder, and adrenal gland(Zou et.al.2020;Qi F et. al. 2020). A varied group of severity and its associated comorbidity upon SARS-CoV-2 infection seems the outcome of differential immune responses due to the differential expressional status of ACE2 under different pathological conditions.

Viral entry depends upon binding of viral spike (S) proteins with cellular ACE2 receptors and primed by host proteases. The recent cellular study also demonstrates that serine protease TMPRSS2 may act as a sound target to inhibit the viral entry to the host cell and may neutralize the virus. Apart from that, ACE2 also postulated for its inflammatory immune network with potential mediators like TLRs, Chemokines, and cytokines in several viral as well as a complex disease. In this context, we tried to buildup interactome network model in Cytoscape to understand the interaction, localization, and co-expression pathway prediction with TLR8, IFNγ, CCR2, ACE2, CCR5, TLR3, TLR7, TNFα to understand the response to the virus, inflammatory response, positive regulation of cytokine production, cell chemotaxis and regulation of inflammatory response. (**Fig 5**).

**Figure 5:**
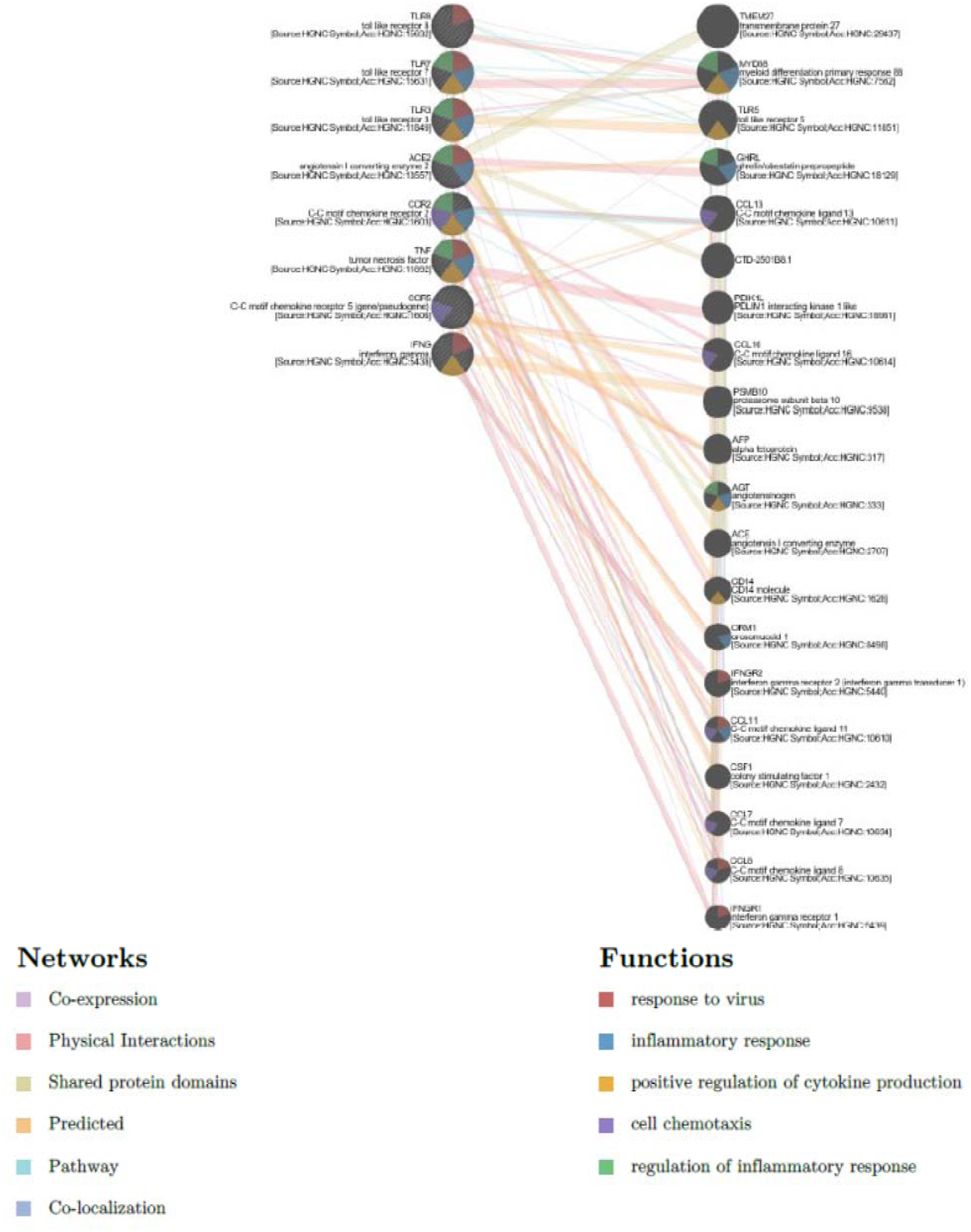
Interactome network model in Cytoscape to understand the interaction, localization, and co-expression with TLR8, IFNγ, CCR2, ACE2, CCR5, TLR3, TLR7, TNFα with SARS-CoV-2.

We may, therefore, conclude that a detailed molecular analysis of the SARS-CoV-2 Genome will provide insight into its origin, evolution, genetic drift as well as potential antigenicity, which can be exploited to develop an effective vaccine. Integrating in-silico based data with available computational tools, we identified several potential antigenic epitopes in the SARS-CoV-2 isolates from India that have the potential to serve as an excellent vaccine candidate.

## Contributions

RD, SK, AYS, SAM, and SP performed in-silico data analysis. SK conceptualized the project and overseen the entire work. RD, SK, SP, SAM, and AYS drafted the manuscript. AS, KP, IM, TS, SS^1^,KP, SS^2^, RM assisted in writing the manuscript, discussion, and references. We thank JBS for critical discussion

## Acknowledgments and Funding

SK and RD thank ICMR (Indian Council of Medical Research), DBT(Department of Biotechnology, India) of indirect funding through manpower support. RD and KP are DBT BioCare Fellows. SS, IM, TS are CSIR fellows. KP and RM are CSIR fellows. We thank Becos Research Lab for providing computational tools. We thank and express our gratitude to the All India Institute of Medical Sciences(AIIMS), New Delhi, India, for financial supporting AS and SS. We thank the Department of Biochemistry, AIIMS, New Delhi for providing logistics and support to carry on this work.

## TABLES

**Table 1A.**
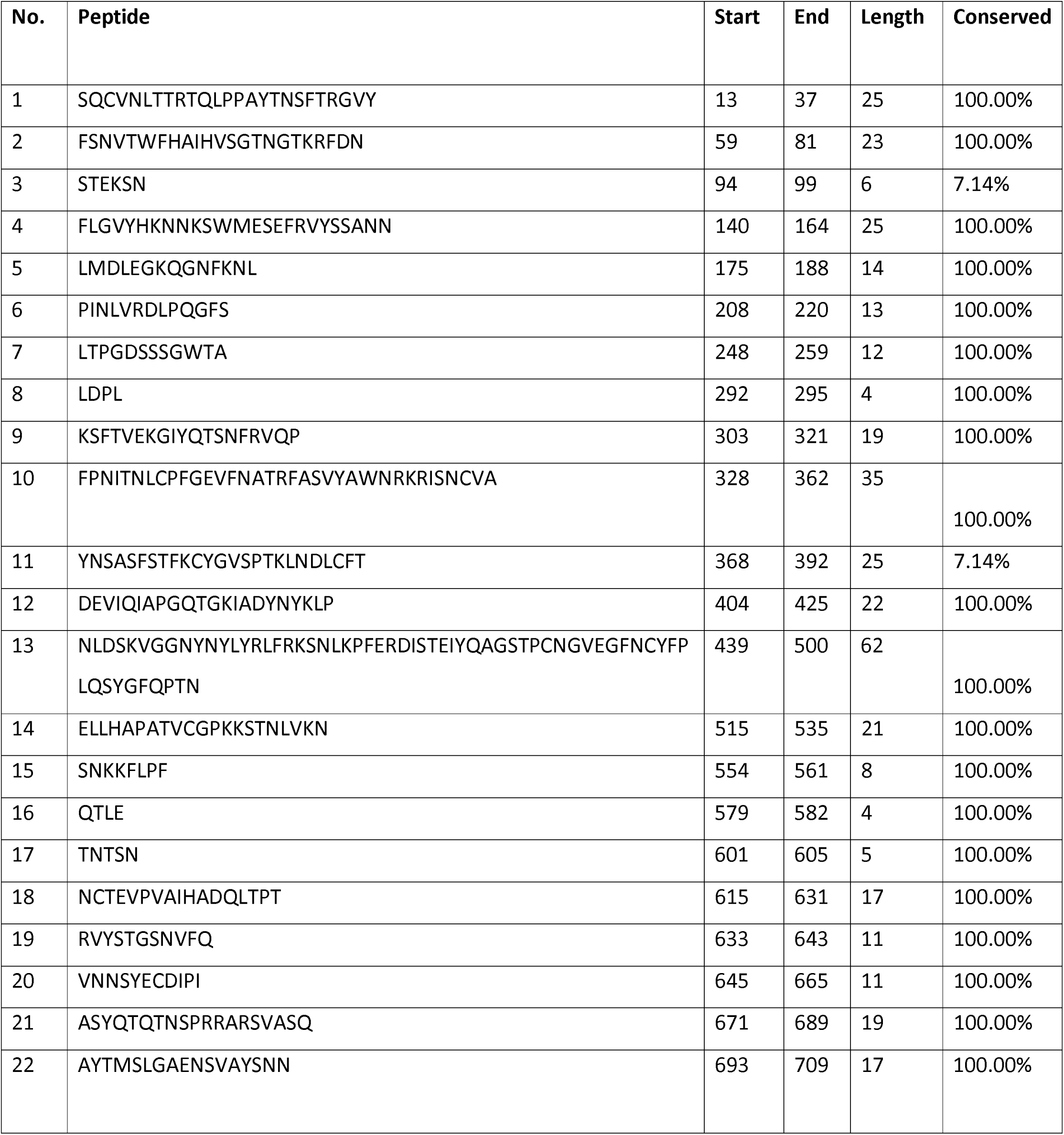
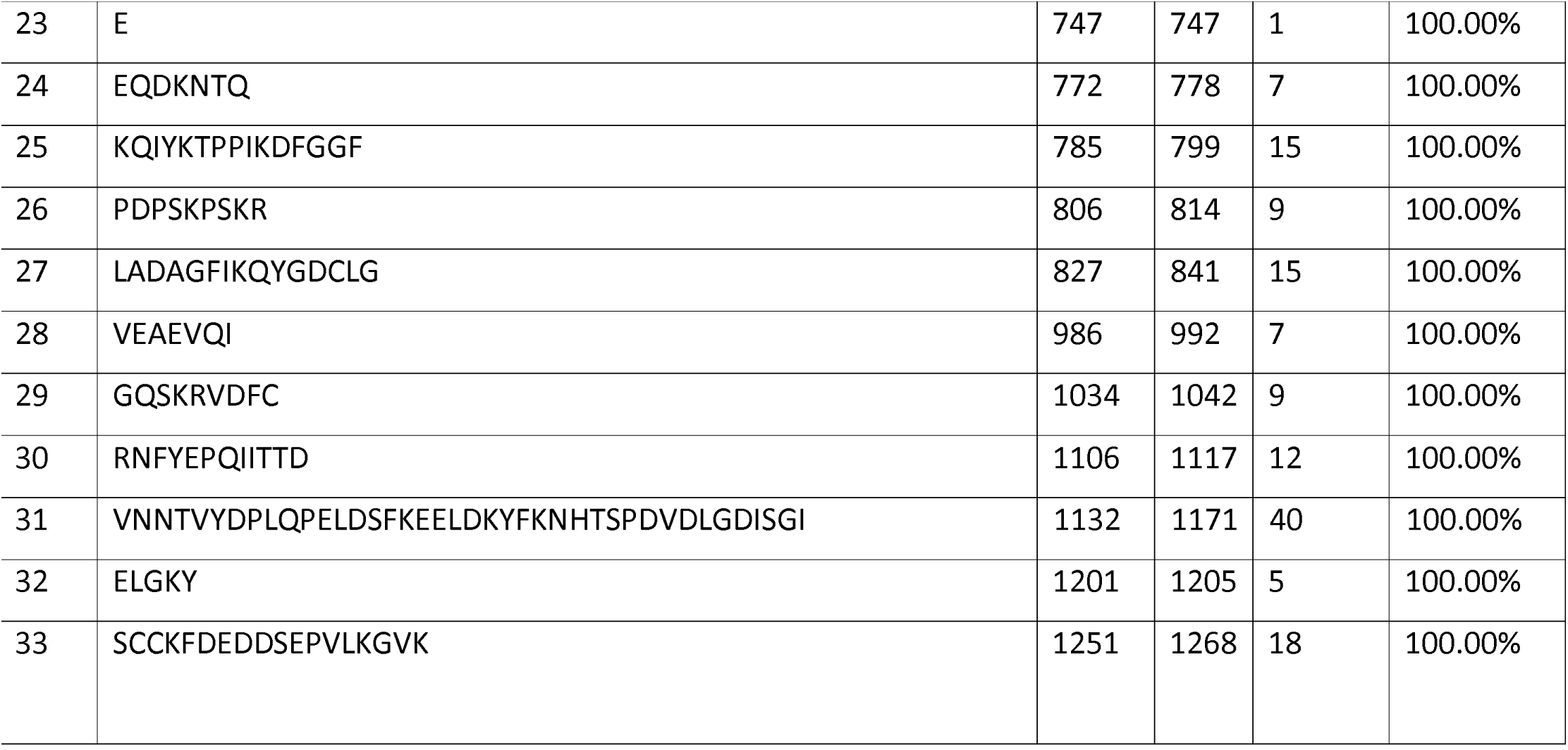
Potential B-cell Linear epitopes (B.E. pipred) for S protein of Indian strain MT012098.

**Table 1B:**
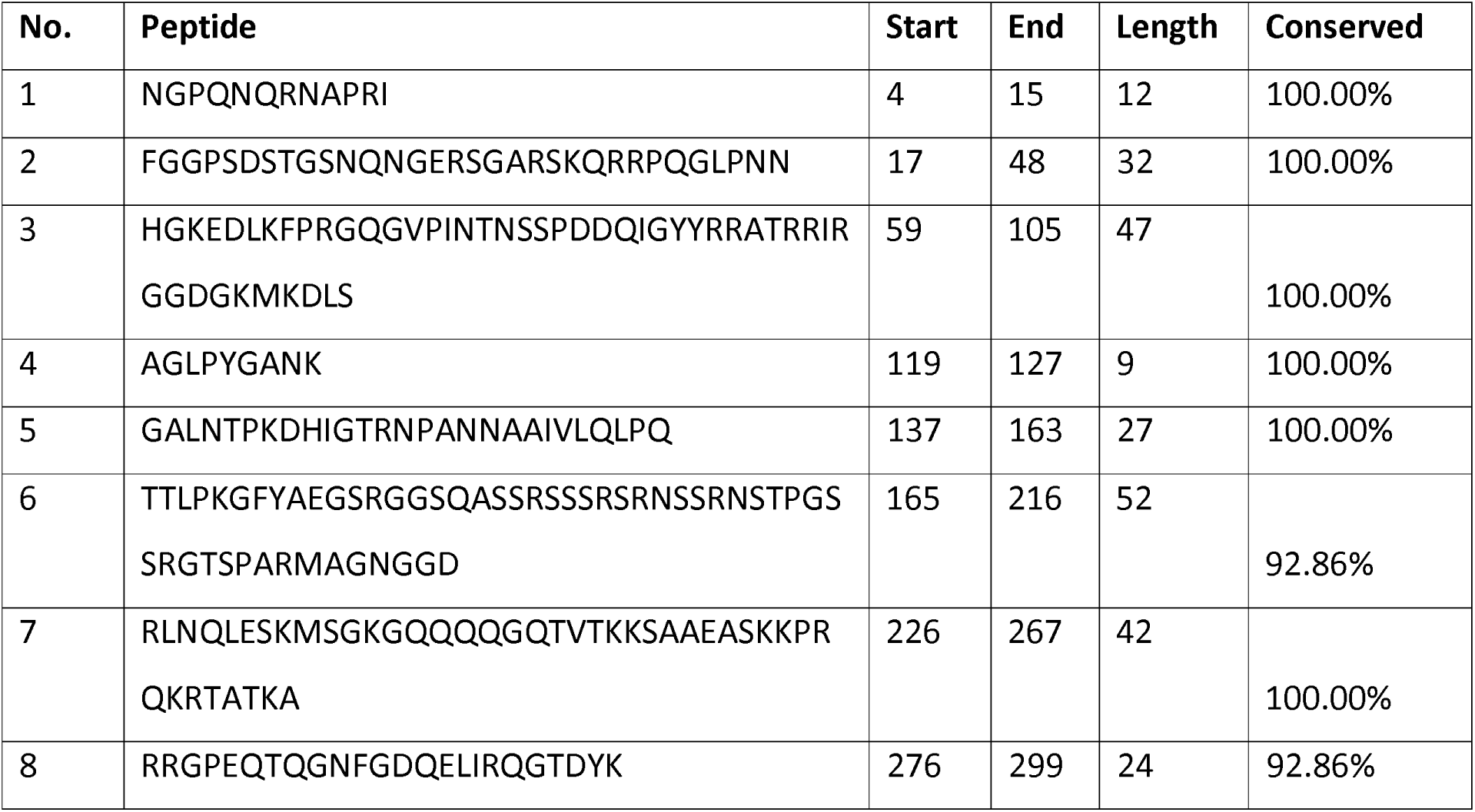
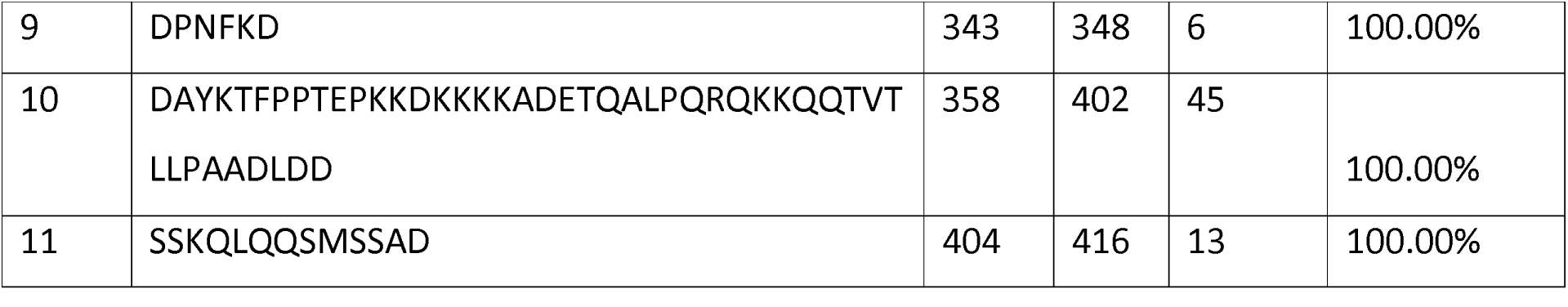
Potential B-cell Linear epitopes (BepiPred) for N protein of Indian strain MT012098.

**Table 1C.** Potential B-cell Linear epitopes (BepiPed) for M protein of Indian strain MT012098.

| No. | Peptide | Start | End | Length | Conserved |
| --- | --- | --- | --- | --- | --- |
| 1 | NGTITVEELKKLLEQW | 5 | 20 | 16 | 100.00% |
| 2 | AN | 40 | 41 | 2 | 100.00% |
| 3 | PLLESE | 132 | 137 | 6 | 100.00% |
| 4 | IKD | 161 | 163 | 3 | 100.00% |
| 5 | KL GASQ RVAGDS | 180 | 191 | 12 | 100.00% |
| 6 | YRIGNYKLNTDHSSSSDNIA | 199 | 218 | 20 | 100.00% |

**Table 1D.**
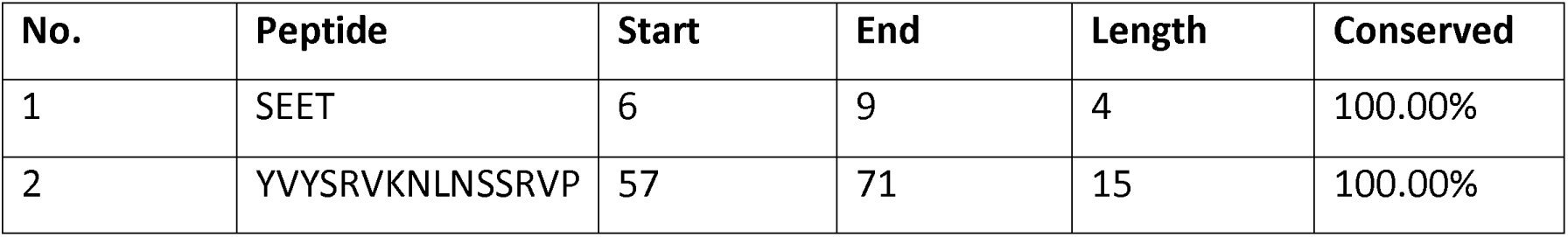
Potential B-cell Linear epitopes (BepiPred) for E protein of Indian strain MT012098.

**Table 1E.**
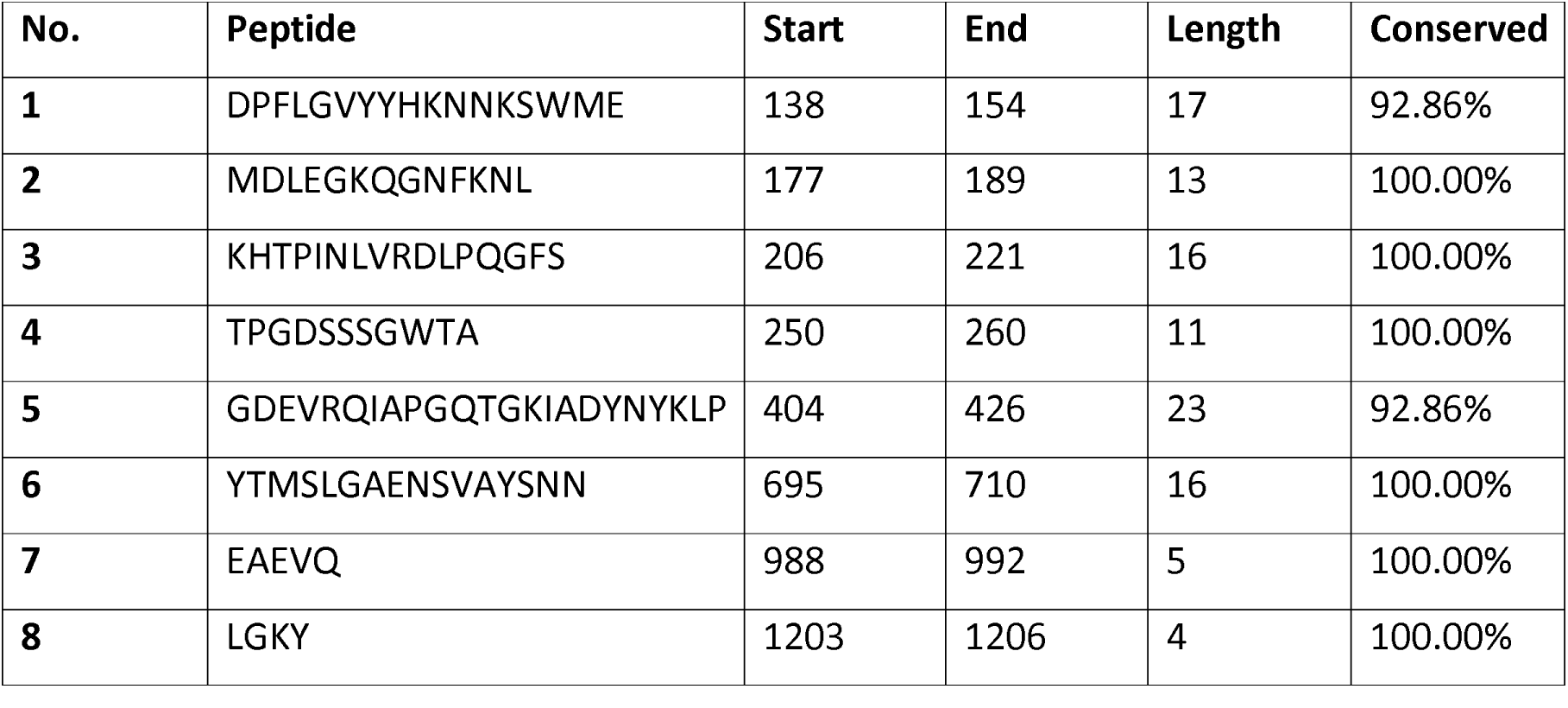
Potential B-cell Linear epitopes (BepiPred) for S protein of reference strain NC_045512 which is not common to the Indian strain MT012098.

**Table 2A.**
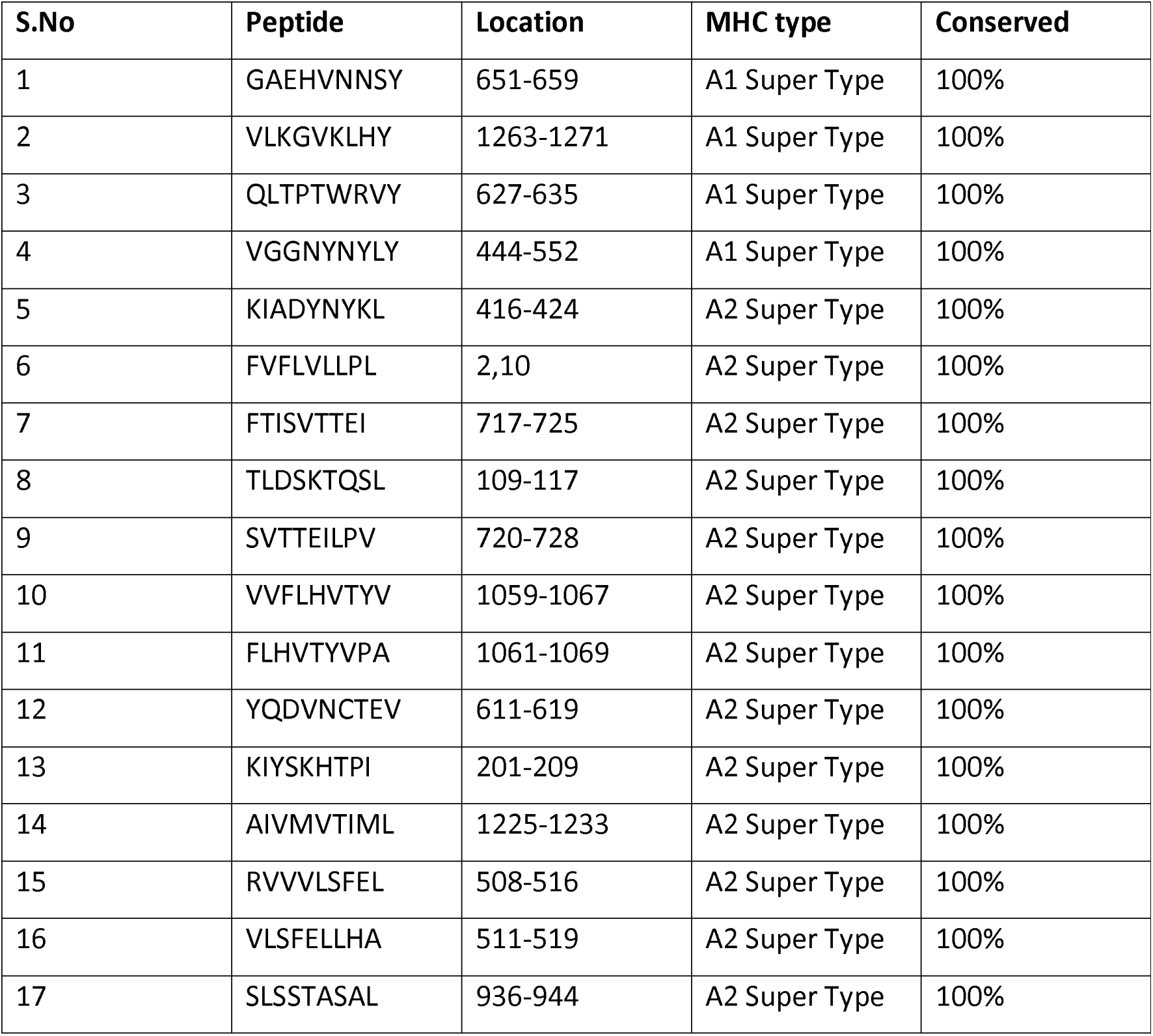
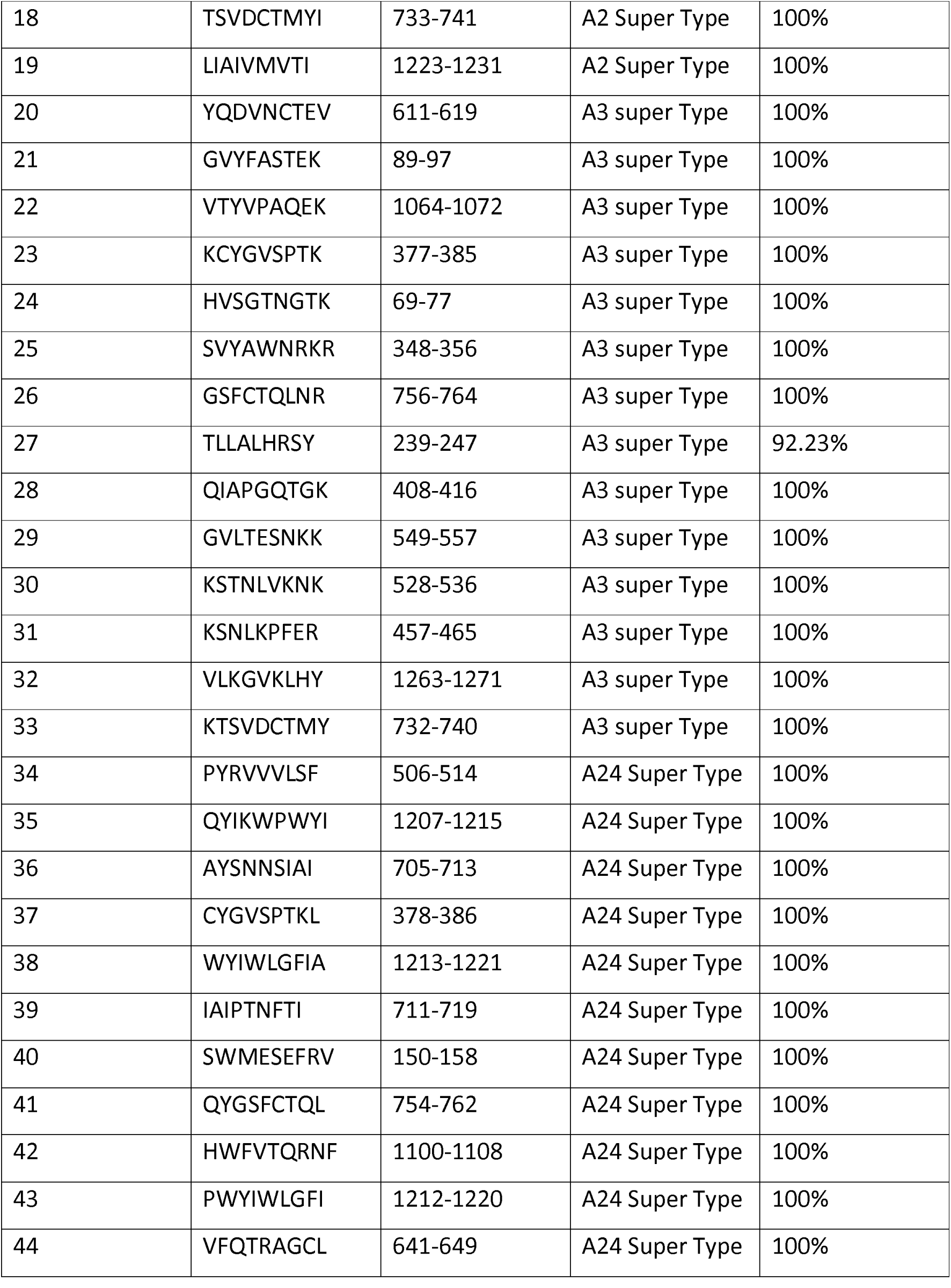
T cell (CTL) Super peptides S protein of Indian strain MT012098 for A1 haplotypes.

**Table 2B.**
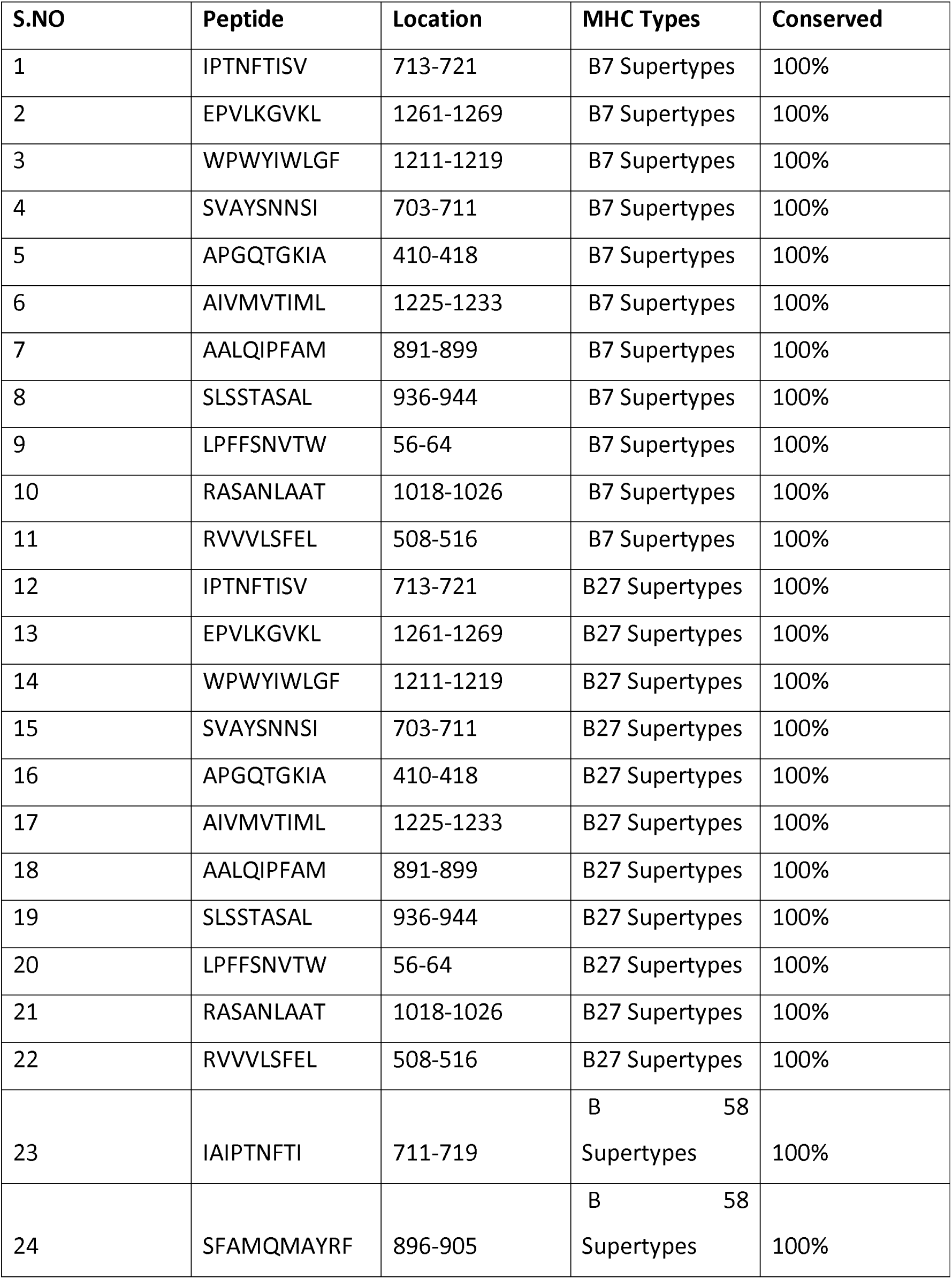
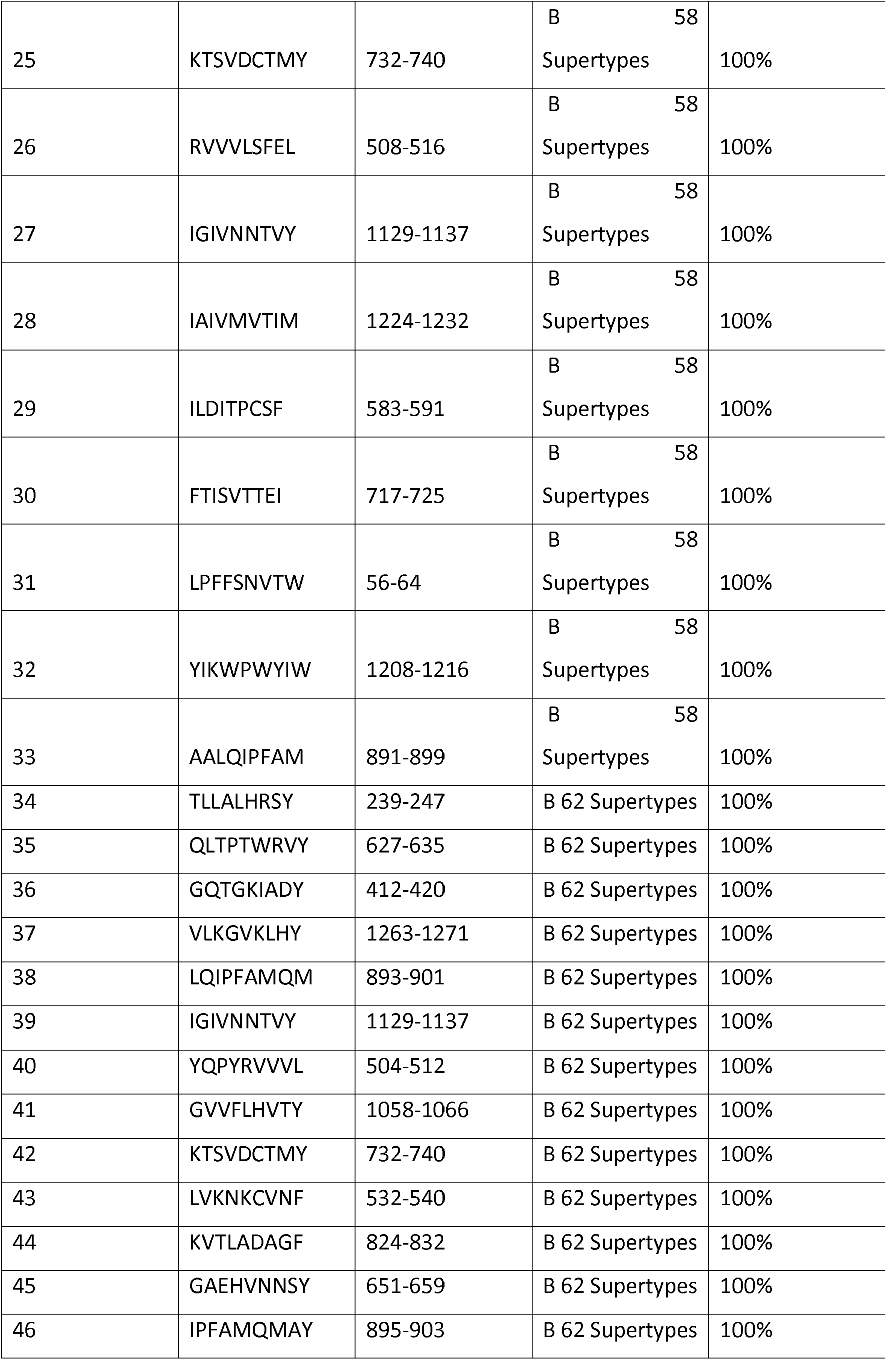
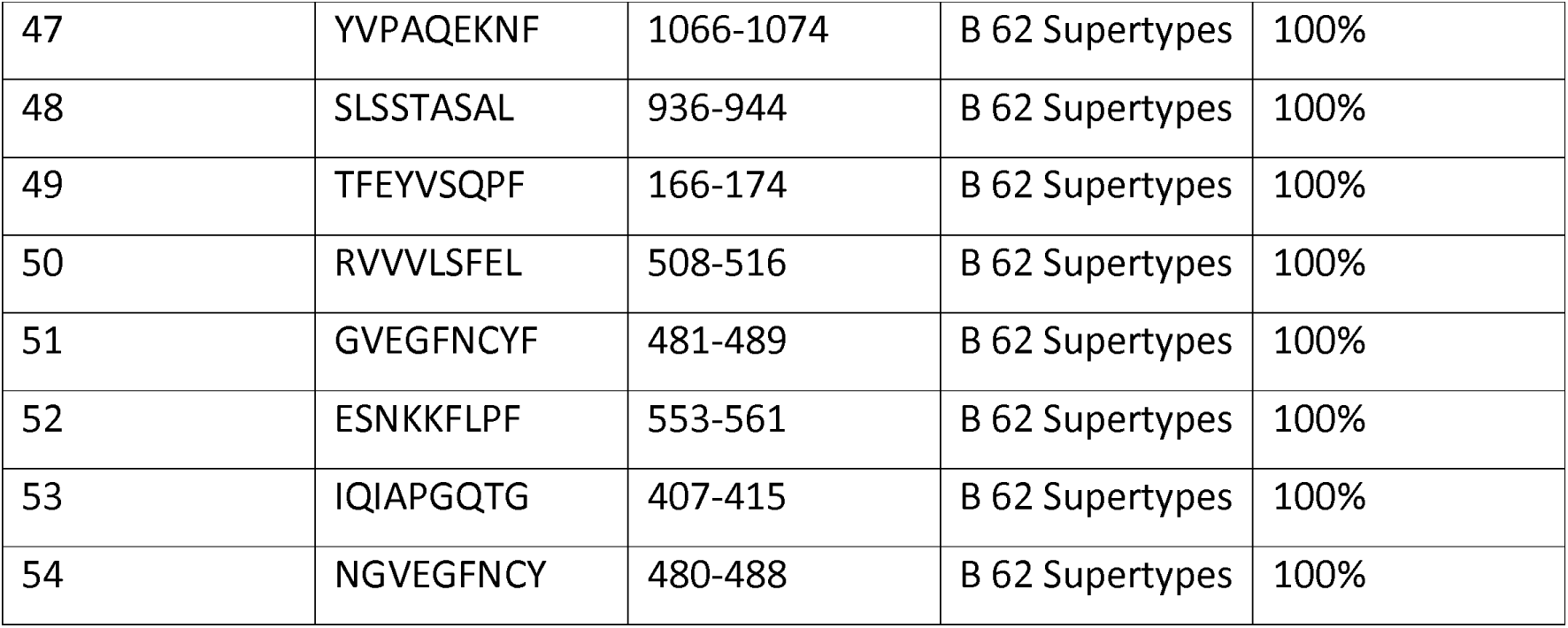
T cell (CTL): Super peptides S protein of Indian strain MT012098 for B1 haplotypes.

**Table 2C.**
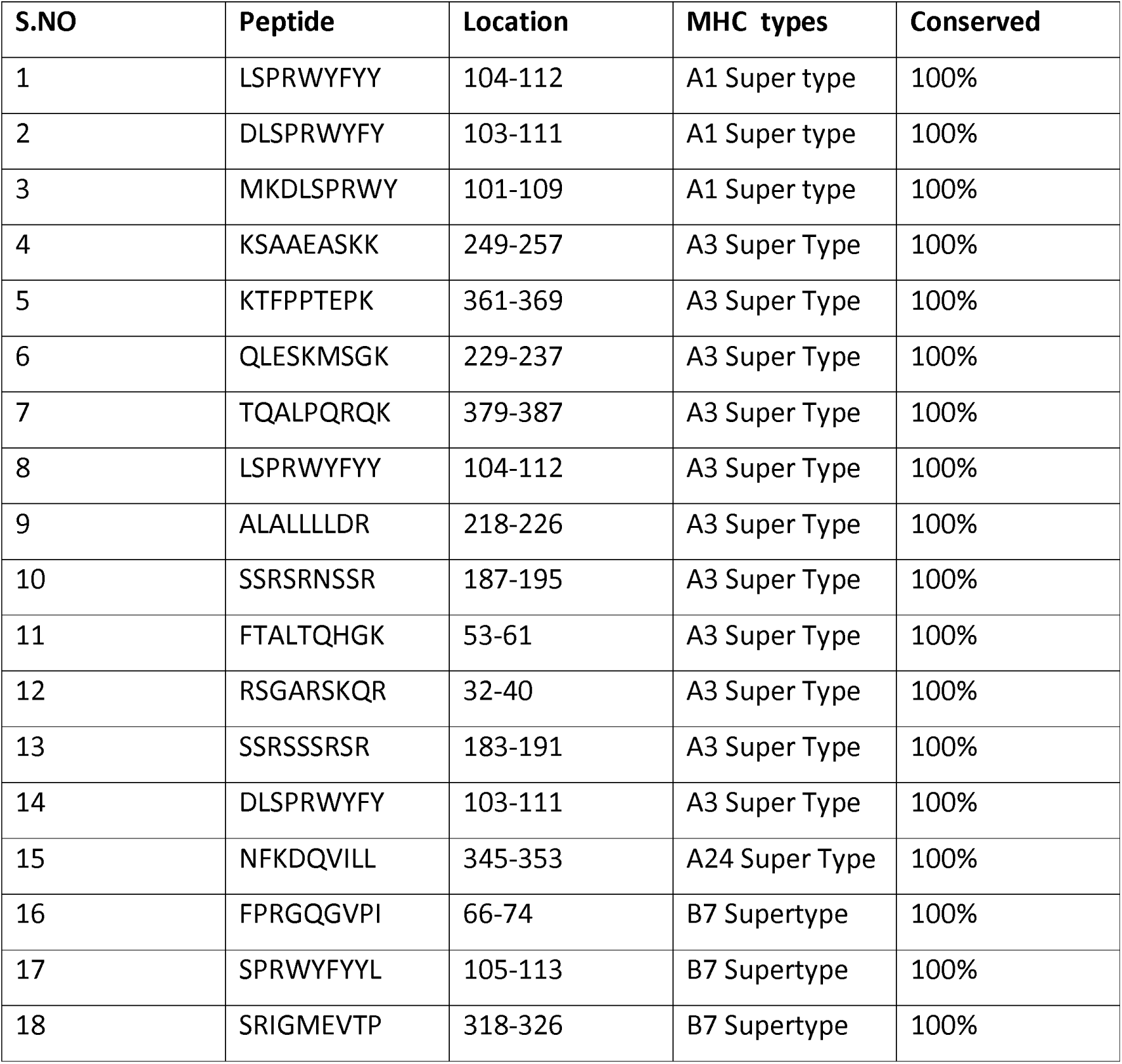
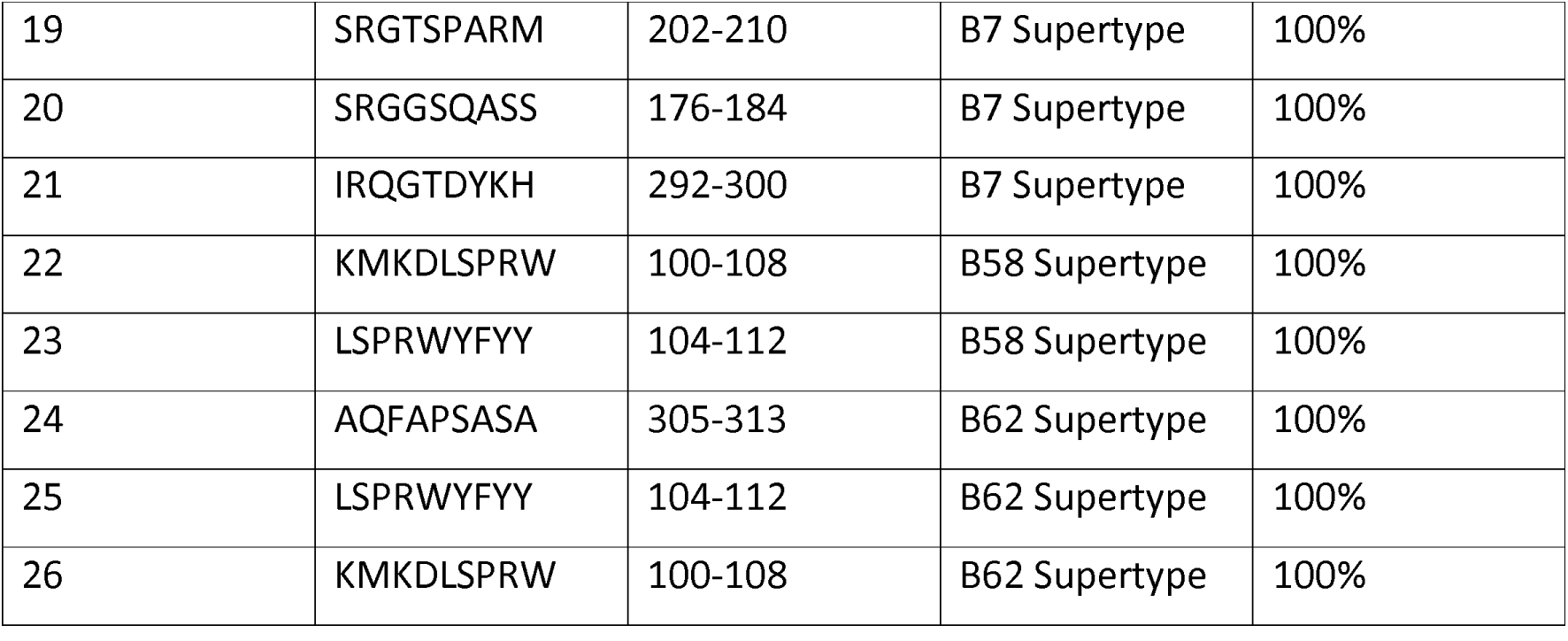
T cell (CTL) Super peptides N protein of Indian strain MT012098.

**Table 2D.**
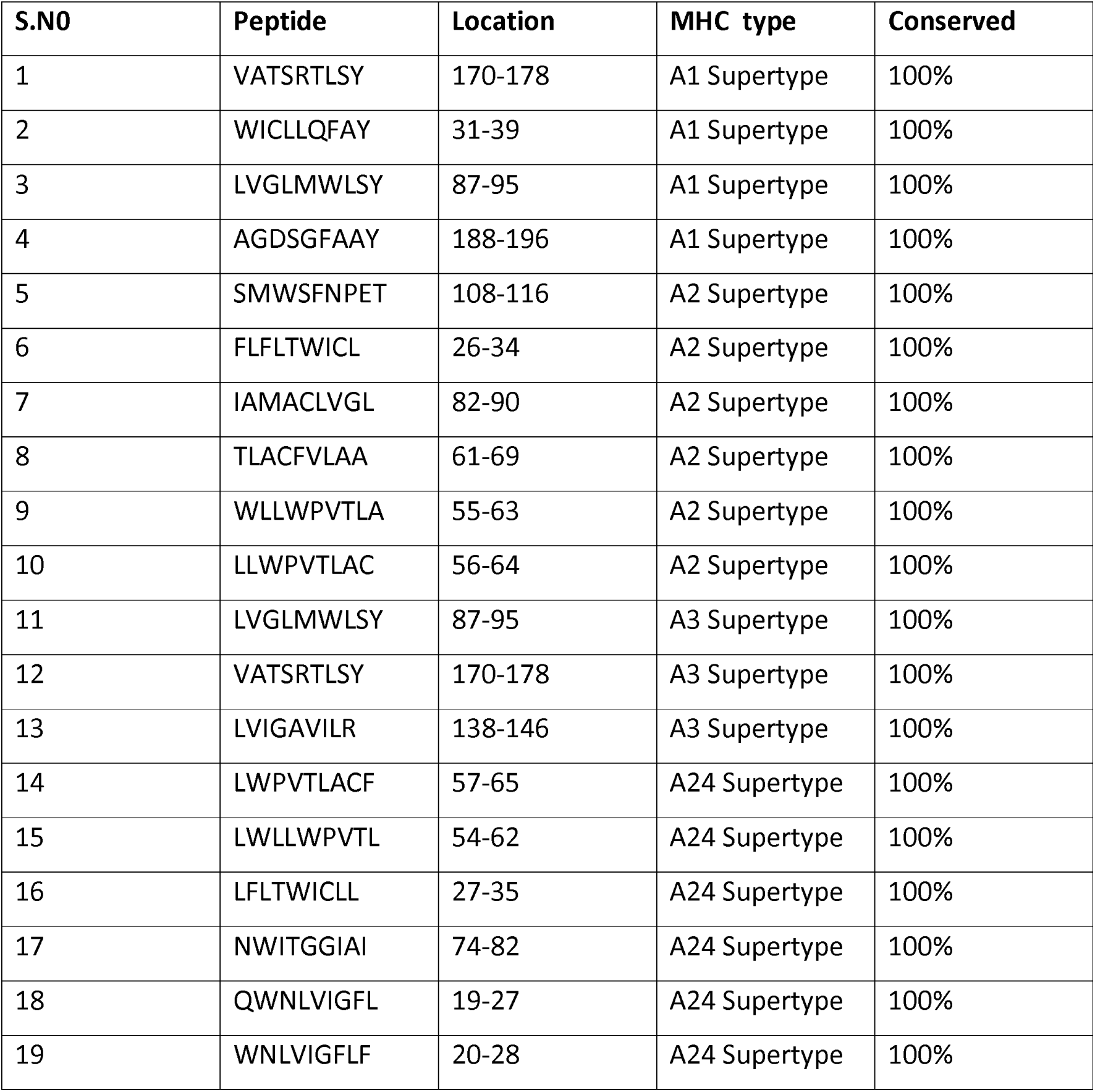
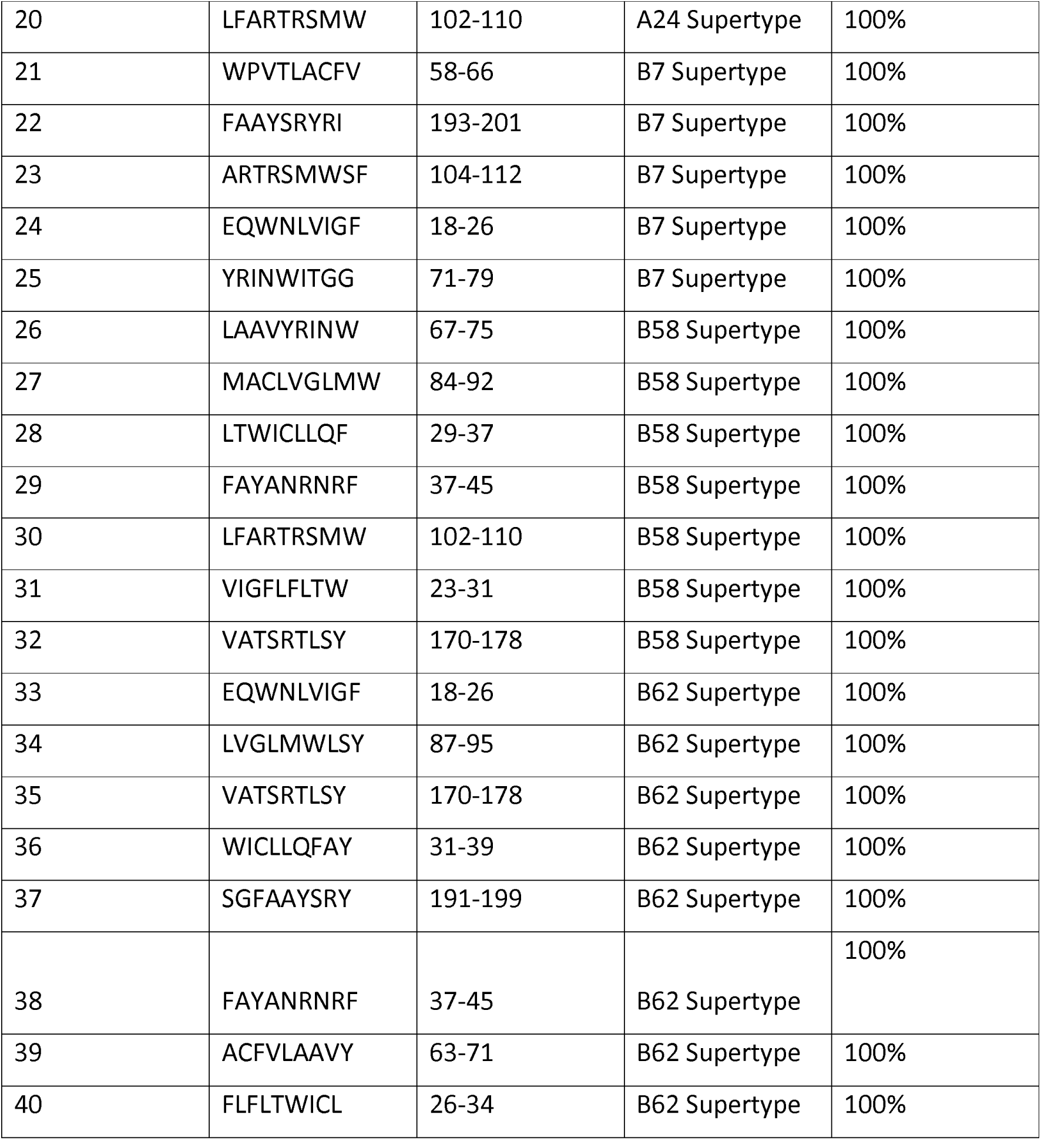
T cell (CTL) Super peptides M protein of Indian strain MT012098.

**Table 2E.**
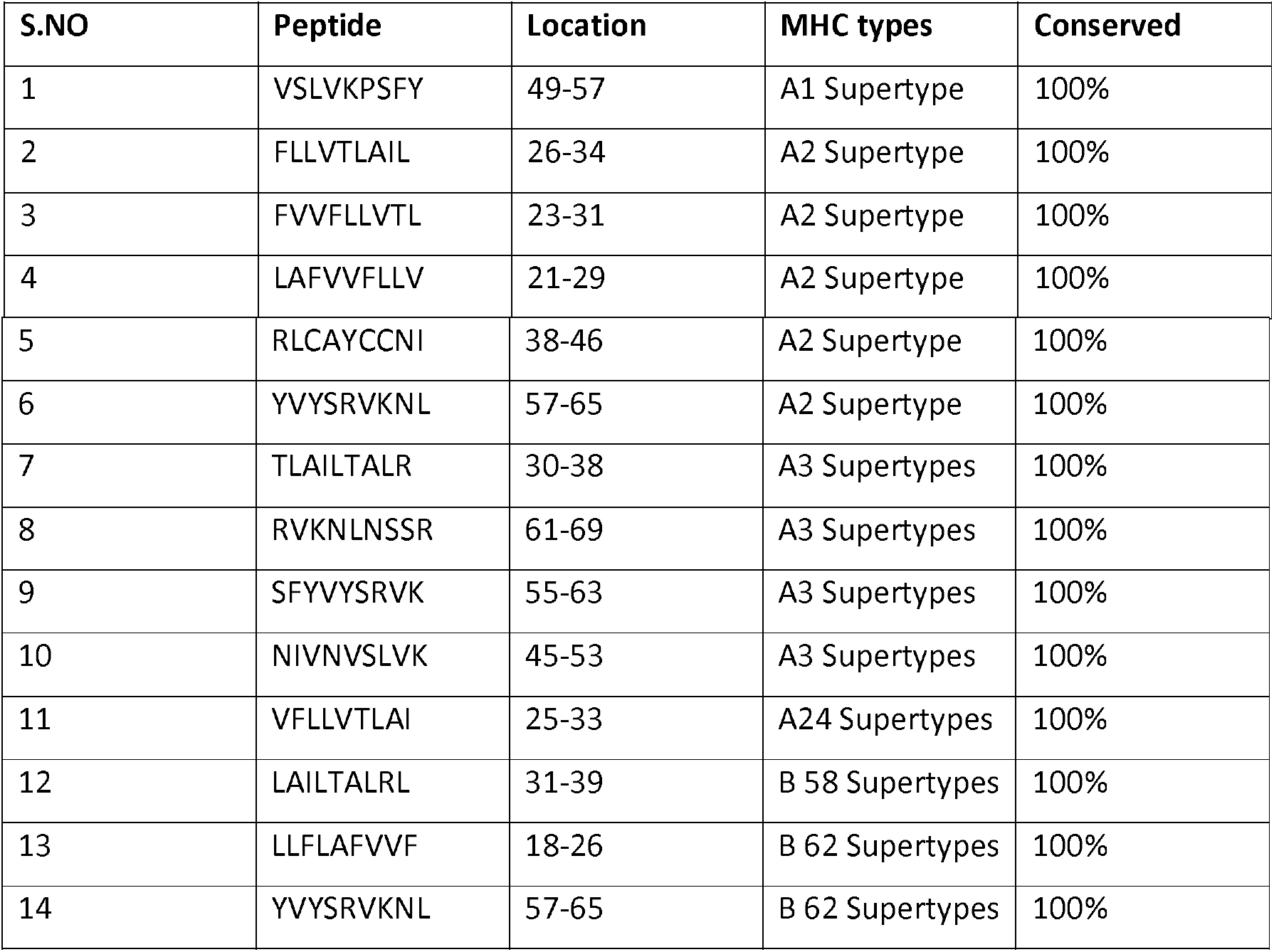
T cell super peptides (CTL) E protein of Indian strain MT012098.

**Table 3A.**
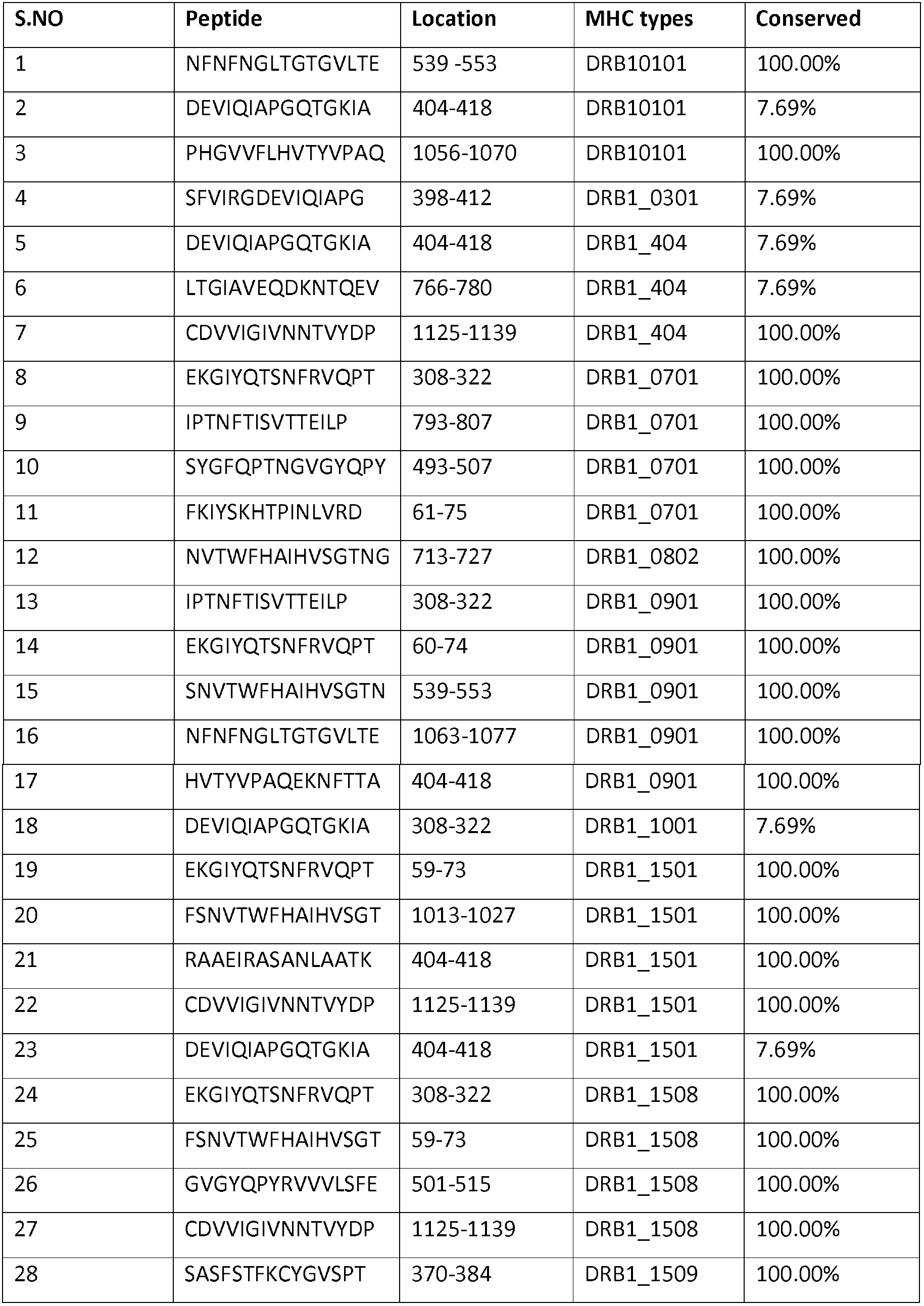
T cell super peptides HLADR CD4^+^ S protein of Indian strain MT012098.

**Table 3B.**
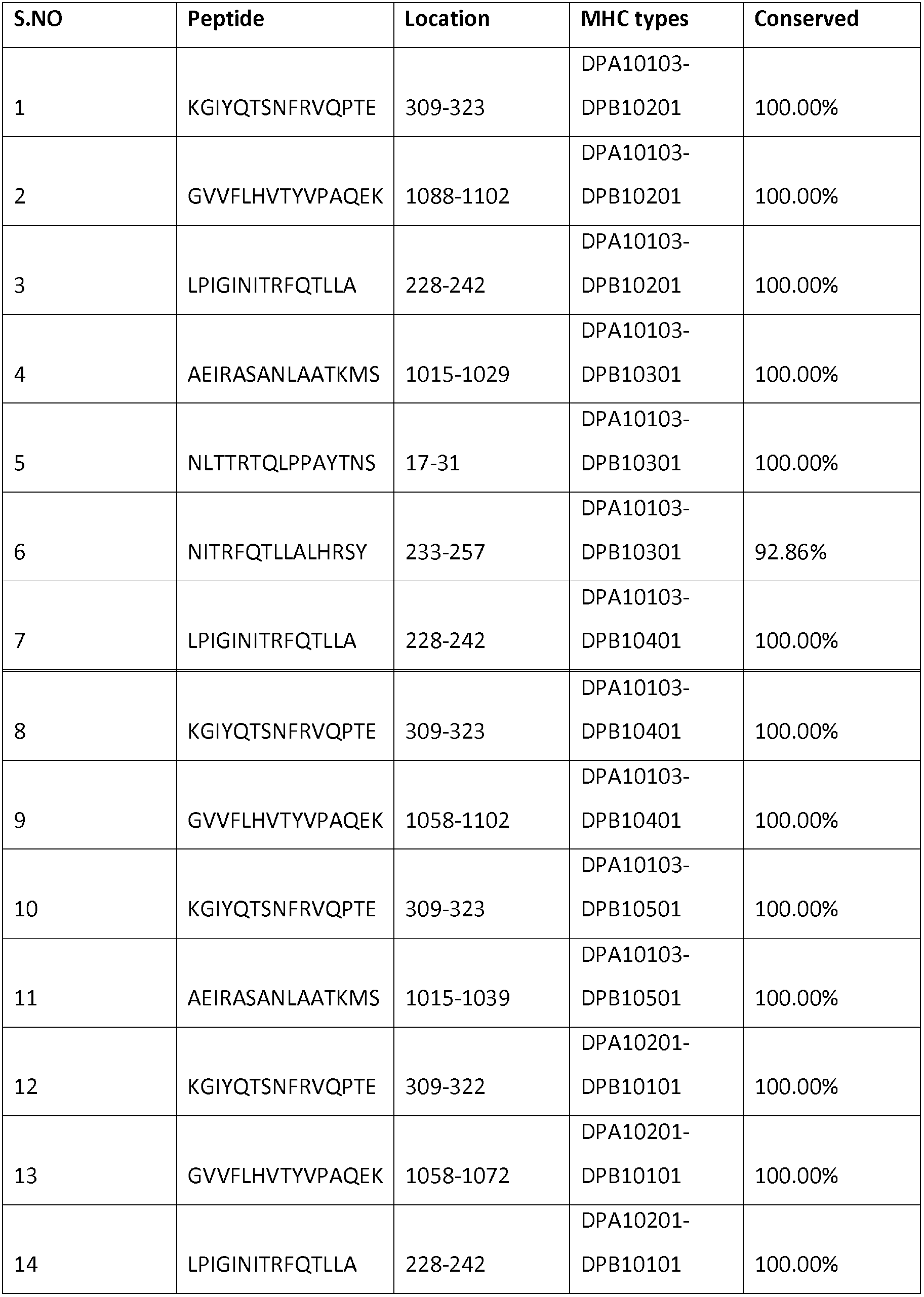
T cell super peptides HLADP CD4^+^S protein of Indian strain MT012098.

**Table 3C.**
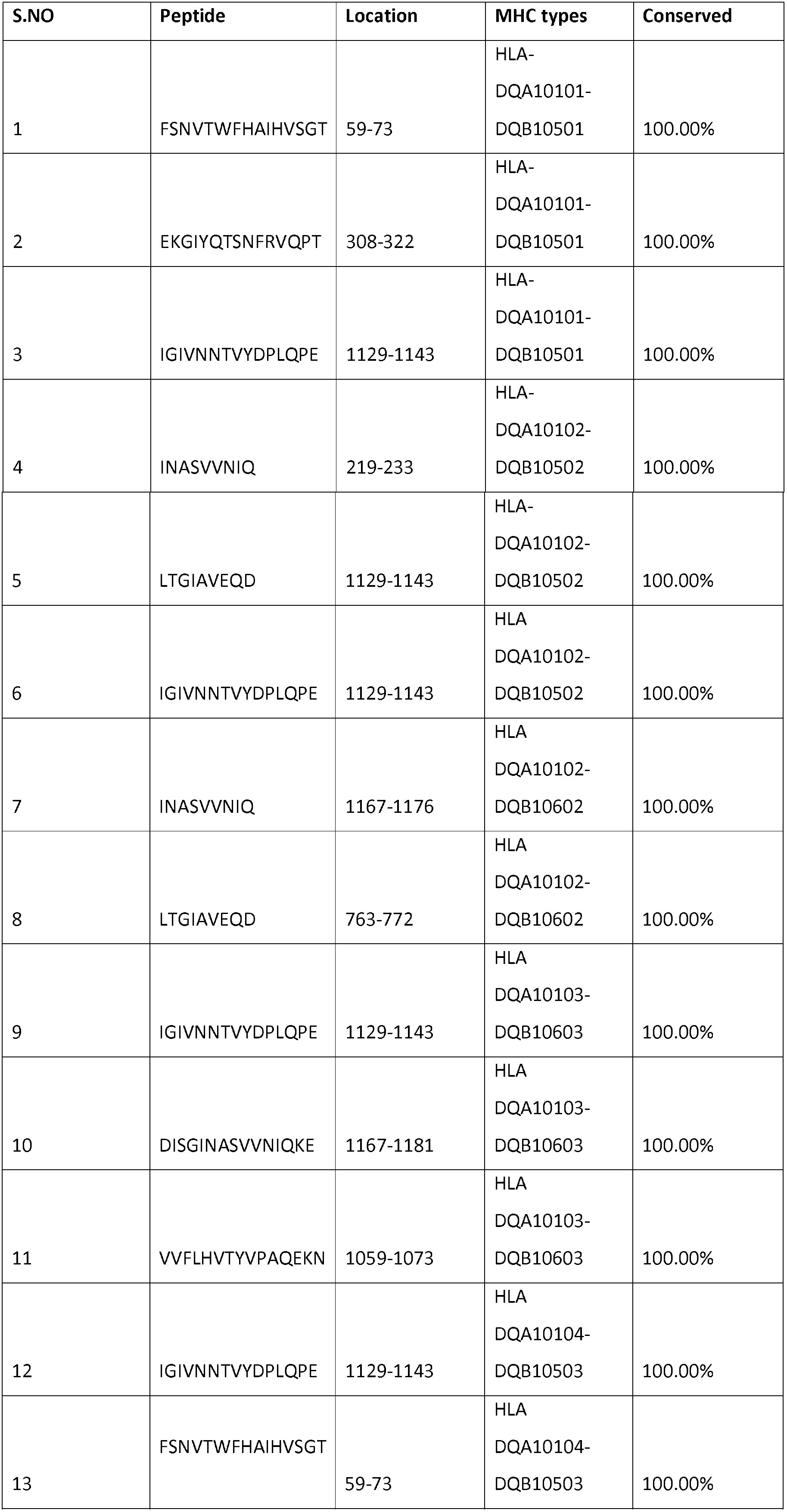
T cell super peptides HLADP CD4^+^S protein of Indian strain MT012098.

**Table 4A.** T cell super peptides HLADP CD4^+^ N protein of Indian strain MT012098.

| S.NO | Peptide | Location | MHC types | Conserved |
| --- | --- | --- | --- | --- |
| 1 | GTWLYTGAIKLDDK | 328-342 | DRB1_0101 | 100.00% |
| 2 | GMSRIGMEVTPSGTW | 316-330 | DRB1_0401 | 100.00% |
| 3 | GTWLYTGAIKLDDK | 328-342 | DRB1_0701 | 100.00% |
| 4 | GTDYKHWPQIAQFAP | 295-309 | DRB1_0802 | 100.00% |
| 5 | GTWLYTGAIKLDDK | 328-342 | DRB1_0901 | 100.00% |
| 6 | KDPNFKDQVILLNKH | 342-356 | DRB1_1001 | 100.00% |
| 7 | GTWLYTGAIKLDDK | 328-342 | DRB1_1508 | 100.00% |
| 8 | GTWLYTGAIKLDDK | 328-342 | DPA10201-DPB10101 | 100.00% |
| 9 | KADETQALPQRQKKQ | 375-389 | DQA10102-DQB10602 | 100.00% |

## References

1. Forster, Peter, et al. “Phylogenetic network analysis of SARS-CoV-2 genomes.” Proceedings of the National Academy of Sciences 117.17 (2020): 9241–9243.

2. Hadfield, James, et al. “Nextstrain: real-time tracking of pathogen evolution.” Bioinformatics 34.23 (2018): 4121–4123.

3. Hatcher, Eneida L., et al. “Virus Variation Resource–improved response to emergent viral outbreaks.” Nucleic acids research 45.D1 (2017): D482–D490.

4. Biswas, Nidhan K., Partha P. Majumder, and West Bengal. “Analysis of RNA Sequences of 3636 SARS-CoV-2 Collected from 55 Countries Reveals Selective Sweep of One Virus Type.” Indian J. Med. Res (2020).

5. Shu, Yuelong, and John McCauley. “GISAID: Global initiative on sharing all influenza data–from vision to reality.” Eurosurveillance 22.13 (2017).

6. Wu, F., Zhao, S. et al A novel coronavirus associated with a respiratory disease in Wuhan of Hubei province, China (Unpublished, preprint)

7. Adhikari, U.K., Tayebi, M., and Rahman, M.M. (2018). Immunoinformatics Approach for Epitope-Based Peptide Vaccine Design and Active Site Prediction against Polyprotein of Emerging Oropouche Virus. J Immunol Res 2018, 6718083.

8. Bui, H.H., Sidney, J., Li, W., Fusseder, N., and Sette, A. (2007). Development of an epitope conservancy analysis tool to facilitate the design of epitope-based diagnostics and vaccines. BMC Bioinformatics 8, 361.

9. Dos Santos Francisco, R., Buhler, S., Nunes, J.M., Bitarello, B.D., Franca, G.S., Meyer, D., and Sanchez-Mazas, A. (2015). HLA supertype variation across populations: new insights into the role of natural selection in the evolution of HLA-A and HLA-B polymorphisms. Immunogenetics 67, 651–663.

10. Doytchinova, I.A., and Flower, D.R. (2007). VaxiJen: a server for prediction of protective antigens, tumour antigens and subunit vaccines. BMC Bioinformatics 8, 4.

11. Jespersen, M.C., Peters, B., Nielsen, M., and Marcatili, P. (2017). BepiPred-2.0: improving sequence-based B-cell epitope prediction using conformational epitopes. Nucleic Acids Res 45, W24–W29.

12. Larsen, M.V., Lundegaard, C., Lamberth, K., Buus, S., Lund, O., and Nielsen, M. (2007). Large-scale validation of methods for cytotoxic T-lymphocyte epitope prediction. BMC Bioinformatics 8, 424

13. Ponomarenko, J., Bui, H.H., Li, W., Fusseder, N., Bourne, P.E., Sette, A., and Peters, B. (2008). ElliPro: a new structure-based tool for the prediction of antibody epitopes. BMC Bioinformatics 9, 514.

14. Thornton JM, Edwards MS, Taylor WR, Barlow DJ. Location of ‘continuous’ antigenic determinants in the protruding regions of proteins. EMBO J. 1986;5(2):409–413.

15. Reynisson, B., Alvarez, B., Paul, S., Peters, B., and Nielsen, M. (2020). NetMHCpan-4.1 and NetMHCIIpan-4.0: improved predictions of MHC antigen presentation by concurrent motif deconvolution and integration of MS MHC eluted ligand data. Nucleic Acids Res.

16. Shankarkumar, U. (2010). Complexities and similarities of HLA antigen distribution in Asian subcontinent. Indian J Hum Genet 16, 108–110

17. Shen, W.J., Zhang, X., Zhang, S., Liu, C., and Cui, W. (2018). The Utility of Supertype Clustering in Prediction for Class II MHC-Peptide Binding. Molecules 23.

18. Sidney, J., Peters, B., Frahm, N., Brander, C., and Sette, A. (2008). HLA class I supertypes: a revised and updated classification. BMC Immunol 9, 1.

19. Zhao, J.W., Yan, M., Shi, G., Zhang, S.L., and Ming, L. (2017). In silico identification of cytotoxic T lymphocyte epitopes encoded by RD5 region of Mycobacterium tuberculosis. J Infect Dev Ctries 11, 806–810.

20. Zhou Y., Fu B., Zheng X., Wnag D., Zhao C., Qi Y., Sun R., Tian Z., Xu X., Wei H. Pathogenic T cells and inflammatory monocytes incite inflammatory storm in severe COVID-19 patients. Journal. 2020

21. Qin C., Zhou L., Hu Z., Zhang S., Yang S., Tao Y., Xie C., Ma K., Shang K., Wang W., Tian D.S. Dysregulation of immune response in patients with COVID-19 in Wuhan, China. Journal. 2020 doi: 10.1093/cid/ciaa248

22. Fu, Y., Cheng, Y. & Wu, Y. Understanding SARS-CoV-2-Mediated Inflammatory Responses: From Mechanisms to Potential Therapeutic Tools. Virol. Sin. (2020). https://doi.org/10.1007/s12250-020-00207-4

23. Yang M (2020) Cell pyroptosis, a potential pathogenic mechanism of 2019-nCoV infection. SSRN. https://doi.org/10.2139/ssrn.3527420

24. Gu J, Gong E, Zhang B, Zheng J, Gao Z, Zhong Y, Zou W, Zhan J, Wang S, Xie Z, Zhuang H, Wu B, Zhong H, Shao H, Fang W, Gao D, Pei F, Li X, He Z, Xu D, Shi X, Anderson VM, Leong AS. Multiple organ infection and the pathogenesis of SARS. J Exp Med. 2005 Aug 1;202(3):415–24. doi: 10.1084/jem.20050828. Epub 2005 Jul 25. PMID: 16043521; PMCID: PMC2213088.

25. Frieman M, Heise M, Baric R. SARS coronavirus and innate immunity. Virus Res. 2008 Apr;133(1):101–12. doi: 10.1016/j.virusres.2007.03.015. Epub 2007 Apr 23. PMID: 17451827; PMCID: PMC2292640.

26. Tay, M.Z., Poh, C.M., Rénia, L. et al. The trinity of COVID-19: immunity, inflammation and intervention. Nat Rev Immunol 20, 363–374 (2020). https://doi.org/10.1038/s41577-020-0311-8

27. Astuti I, Ysrafil. Severe Acute Respiratory Syndrome Coronavirus 2 (SARS-CoV-2): An overview of viral structure and host response. Diabetes Metab Syndr. 2020 Apr 18;14(4):407–412. doi: 10.1016/j.dsx.2020.04.020. Epub ahead of print. PMID: 32335367; PMCID: PMC7165108.

28. Li, G, Fan, Y, Lai, Y, et al. Coronavirus infections and immune responses. J Med Virol. 2020; 92: 424–432. https://doi.org/10.1002/jmv.25685

29. Kawai T, Akira S. The role of pattern-recognition receptors in innate immunity: update on Toll-like receptors. Nature Immunol. 2010; 11(5): 373–384.

30. Pobezinskaya YL, Kim YS, Choksi S, et al. The function of TRADD in signaling through tumor necrosis factor receptor 1 and TRIF-dependent Toll-like receptors. Nature Immunol. 2008; 9(9): 1047–1054.

31. Ermolaeva MA, Michallet MC, Papadopoulou N, et al. Function of TRADD in tumor necrosis factor receptor 1 signaling and in TRIF-dependent inflammatory responses. Nature Immunol. 2008; 9(9): 1037–1046.

32. Zheng C, Chen J, Chu F, Zhu J, Jin T. Inflammatory Role of TLR-MyD88 Signaling in Multiple Sclerosis. Front Mol Neurosci. 2020;12:314. Published 2020 Jan 10. doi:10.3389/fnmol.2019.00314

33. Perales-Linares, R. and Navas-Martin, S. (2013), Toll-like receptor 3 in viral pathogenesis: friend or foe?. Immunology, 140: 153–167. doi:10.1111/imm.12143

34. Nilsen NJ, Vladimer GI, Stenvik J, et al. A role for the adaptor proteins TRAM and TRIF in toll-like receptor 2 signaling. J Biol Chem. 2015;290(6):3209–3222. doi:10.1074/jbc.M114.593426.

35. Glass WG, Rosenberg HF, Murphy PM. Chemokine regulation of inflammation during acute viral infection. Curr Opin Allergy Clin Immunol. 2003;3(6):467–473.

36. van Meer, G., Voelker, D. & Feigenson, G. Membrane lipids: where they are and how they behave. Nat Rev Mol Cell Biol 9, 112–124 (2008). https://doi.org/10.1038/nrm2330

37. Gay, N., Symmons, M., Gangloff, M. et al. Assembly and localization of Toll-like receptor signalling complexes. Nat Rev Immunol 14, 546–558 (2014). https://doi.org/10.1038/nri3713

38. Shi-Hui Sun, Qi Chen, Hong-Jing Gu, Guan Yang, Yan-Xiao Wang, Xing-Yao Huang, Su-Su Liu, Na-Na Zhang, Xiao-Feng Li, Rui Xiong, Yan Guo, Yong-Qiang Deng, Wei-Jin Huang, Quan Liu, Quan-Ming Liu, Yue-Lei Shen, Yong Zhou, Xiao Yang, Tong-Yan Zhao, Chang-Fa Fan, Yu-Sen Zhou, Cheng-Feng Qin, You-Chun Wang. A mouse model of SARS-CoV-2 infection and pathogenesis. Cell Host & Microbe, 2020; DOI: 10.1016/j.chom.2020.05.020

39. Jason Netland, David, K. Meyerholz, Steven Moore, Martin Cassell, Stanley Perlman. Journal of Virology Jul 2008, 82 (15) 7264–7275; DOI: 10.1128/JVI.00737-08

40. Coperchini F, Chiovato L, Croce L, Magri F, Rotondi M. The cytokine storm in COVID-19: An overview of the involvement of the chemokine/chemokine-receptor system [published online ahead of print, 2020 May 11]. Cytokine Growth Factor Rev. 2020;S1359-6101(20)30092-7. doi:10.1016/j.cytogfr.2020.05.003

41. Zhang C, Wu Z, Li JW, Zhao H, Wang GQ. Cytokine release syndrome in severe COVID-19: interleukin-6 receptor antagonist tocilizumab may be the key to reduce mortality. Int J Antimicrob Agents. 2020;55(5):105954. doi:10.1016/j.ijantimicag.2020.105954

42. Bourgonje AR, Abdulle AE, Timens W, et al. Angiotensin-converting enzyme-2 (ACE2), SARS-CoV-2 and pathophysiology of coronavirus disease 2019 (COVID-19) [published online ahead of print, 2020 May 17]. J Pathol. 2020; 10.1002/path.5471. doi:10.1002/path.5471

43. Zou X, Chen K, Zou J, Han P, Hao J, Han Z. Single-cell RNA-seq data analysis on the receptor ACE2 expression reveals the potential risk of different human organs vulnerable to 2019-nCoV infection. Front Med. 2020;14(2):185–192. doi:10.1007/s11684-020-0754-0

44. Qi F, Qian S, Zhang S, Zhang Z. Single cell RNA sequencing of 13 human tissues identify cell types and receptors of human coronaviruses. Biochem Biophys Res Commun. 2020;526(1):135–140. doi:10.1016/j.bbrc.2020.03.044

